# Extracellular vesicles from microglial cells activated by abnormal heparan sulfate oligosaccharides from Sanfilippo patients impair neuronal dendritic arborization

**DOI:** 10.1101/2024.05.22.595318

**Authors:** Chloé Dias, Nissrine Ballout, Guillaume Morla, Katia Alileche, Christophe Santiago, Chiara Ida Guerrera, Adeline Chaubet, Jerome Ausseil, Stephanie Trudel

## Abstract

**Background:** In mucopolysaccharidosis type III (MPS III), a pediatric neurodegenerative disorder, accumulation of abnormal glycosaminoglycans (GAGs) induces severe neuroinflammation by triggering the microglial pro-inflammatory cytokines production via a TLR4-dependent pathway. But the extent of the microglia contribution to the MPS III neuropathology remains unclear. Extracellular vesicles (EVs) mediate intercellular communication and are known to participate in the pathogenesis of adult neurodegenerative diseases. However, characterization of the molecular profiles of EVs released by MPS III microglia and their effects on neuronal functions have not been described.

**Methods:** Here, we isolated EVs secreted by the microglial cells after treatment with GAGs purified from urines of MPS III patients (MPS III-EVs) to explore the EVs’ proteins and small RNA profiles using LC-MS/MS and RNA sequencing. We next performed a functional assay by immunofluorescence following wild-type (WT) or MPS III-EVs uptake by WT primary cortical neurons and analyzed their extensions metrics after staining of βIII-tubulin and MAP2 by confocal microscopy.

**Results:** Functional enrichment analysis for both proteomics and RNA sequencing data from MPS III-EVs revealed a specific content involved in neuroinflammation and neurodevelopment pathways. Treatment of cortical neurons with MPS III-EVs induced a disease-associated phenotype demonstrated by a lower total neurite surface area, an impaired somatodendritic compartment, and a higher number of immature dendritic spines.

**Conclusions:** This study shows, for the first time, that GAGs from patients with MPS III can induce microglial secretion of EVs that deliver a specific molecular message to recipient naive neurons, while promoting the neuroinflammation, and depriving neurons of neurodevelopmental factors. This work provides a framework for further studies of biomarkers to evaluate efficiency of emerging therapies.

**Graphical abstract:** 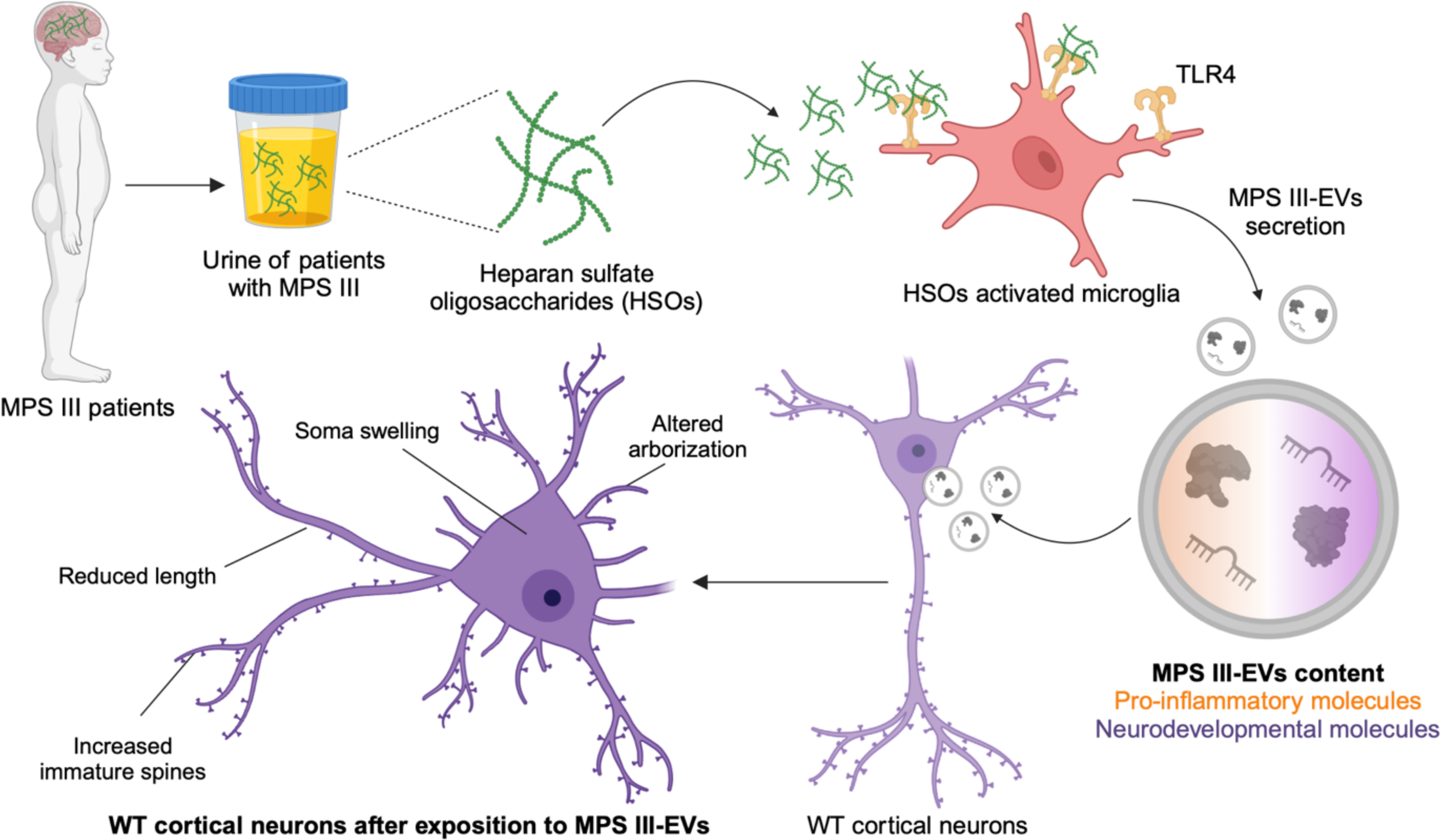

## Background

Mucopolysaccharidosis type III (MPS III, also known as Sanfilippo syndrome) is a lysosomal storage disease (LSD) caused by the specific inability to degrade the glycosaminoglycan (GAG) heparan sulfate (HS) – an essential component of the cell surface and the extracellular matrix. There are four subtypes of MPS III (A, B, C and D), depending on the mutated gene and the resulting enzyme deficiency (1). There are no disease-modifying treatments for this rare but ultimately fatal disorder (incidence: ∼1 per 70.000 live births) (2). A number of innovative therapies for MPS III are being studied, and some reaching the mid-to-late stages of clinical development (3), like gene therapy in MPSIIIB patients (4,5). However, there is still a lack of neuro-cognitive response-to-treatment biomarkers and robust blood and CSF-based biomarkers are an urgent unmet need to evaluate therapies efficacy (6). The partially degraded heparan sulfate oligosaccharides (HSOs) accumulate not only in the lysosomal environment but also in other subcellular locations, the extracellular space, tissues, and fluids (7) (including urine (8–10)). In MPS III, the accumulation of highly sulfated and acetylated HSOs in brain tissues causes neuronal dysfunction and neuronal death; these ultimately lead to neuropsychiatric problems, developmental delays, childhood dementia, blindness, and death in the second decade of life (11). Interestingly, HSO accumulation is associated with the progressive aggregation of Aβ, tau, α-synuclein, and prion proteins in brain tissue in young children with MPS III (12–14). This reflects the pathophysiology of adult neurodegenerative diseases because heparan-sulfate proteoglycans (HSPGs) and HS are also found in the amyloid plaques in Alzheimer’s disease, Down’s syndrome, and prion diseases (15). Moreover, a link between MPS III and Parkinson’s disease was suggested when mutations causing the former were linked to a higher risk of developing the latter (16). Even though adult neurodegenerative diseases and MPS III have different molecular and cellular pathogenic mechanisms, chronic neuroinflammation is a hallmark of both conditions.

We have demonstrated previously that the progressive accumulation of abnormal HSOs leads to the Toll-like receptor (TLR) 4-dependent activation of microglia through the MyD88 adaptor and the NF-kB pathway (10,17), which in turn results in neurodegeneration. Strong activation of resident microglia and the secretion of high levels of inflammatory cytokines were subsequently described early in the course of disease in canine and murine models of different subtypes of MPS III (6,10,18–21). Higher expression levels of proteins related to immunity, macrophage function and oxidative stress have been reported in the animal model’s brain tissues and in human cerebrospinal fluid and plasma samples (22–24). Although microglia-mediated inflammation is a key player in the pathogenesis of progressive neurological diseases, the mechanisms by which microglia influence the onset and propagation of neuroinflammation and neurodegeneration remain unclear.

Several converging lines of evidence indicate that extracellular vesicles (EVs) secreted by microglia have an essential role in not only the propagation of misfolded proteins such as tau and ⍺-synuclein (25–27) but also the transmission of inflammation, the impairment of neuronal functions, and the promotion of neurodegeneration (28–31). EVs are lipid-bilayer-delimited particles released by a variety of cells, including neurons, microglia and astrocytes and which carry their specific cargos of proteins, RNAs and lipids over long distances. EV biogenesis involves direct, outward budding from the plasma membrane forming ectosome-like EVs or by inward invagination of the endosomal membrane and fusion of the multivesicular body with the plasma membrane forming exosome-like EVs (32,33).

The contribution of EVs to the pathophysiology – and more specifically to the neuroinflammation – of MPS III or any neuropathic lysosomal storage diseases has not previously been characterized. In the present study, we looked at whether or not MPS III microglia-derived EVs i) can drive a disease-specific content, ii) are involved in the functional impairment of cortical neurons, and iii) can be considered as good and relevant surrogate biomarkers in MPS III. To address these questions, we isolated and characterized EVs released by microglia murine cell line treated with the accumulated substrate at the origin of the neuropathy, GAGs (enriched in HSOs) extracted from the urine of children with MPS III. Omics analyses at the protein and RNA levels revealed that these EVs carried a characteristic cargo to recipient cells in central nervous system (CNS) and were involved in the inflammatory response and neuronal development. After exposing primary cortical neurons isolated from wild-type (WT) pups to MPS III microglia-derived EVs, we used neurite imaging assays to show that the morphology of the somatodendritic compartment was impaired. These findings provide new insights in the comprehension concerning the spreading of neuroinflammation and in the neuronal dysfunction in MPS III, while offering promising perspectives for the development of biomarkers.

## Methods

### Patients, urine collection, and GAG purification

#### Patients with MPS III

All six patients (aged from 6 to 28 years) had a diagnosis of MPS III (A, B or C). They all presented with mental retardation and high concentrations of GAGs in the urine and have no curative treatments. In line with the French Public Health Code (*Code de la Santé Publique*, article L1211-2) approval by an investigational review board was neither required nor sought. However, the parents of under-18 MPSIII patients and/or their legal guardians stated that they did not object to the use of urine samples for biomedical research.

#### Collection of urine samples

Control urine samples from healthy children were obtained from our institution’s biological resource center (Toulouse BioRessources, Toulouse, France). Samples of urine from patients with MPS III patients and from age-matched unaffected donors were collected under sterile conditions and immediately stored at -20°C. The characteristics of patients and controls are described in Table 1.

**Table 1:** Characteristics of urine samples from patients and controls.

| Patients/<br>controls | Clinical<br>type | Sex | Age at diagnosis<br>(months) | Age at the time of urine<br>collection (years) | Mean GAG concentration<br>(mg/L of urine) |
| --- | --- | --- | --- | --- | --- |
| CTL 1 | / | M | / | 15 | 4.9 |
| CTL 2 | / | F | / | 8 | 2.5 |
| CTL 3 | / | F | / | 10 | 5.6 |
| CTL 4 | / | F | / | 6 | 3.9 |
| MPS 1 | MPS IIIA | M | 42 | 12 | 75.3 |
| MPS 2 | MPS IIIB | F | 23 | 6 | 21.1 |
| MPS 3 | MPS IIIC | M | 40 | 7 | 35.2 |
| MPS 4 | MPS IIIC | M | 60 | 17 | 28.1 |
| MPS 5 | MPS IIIC | F | 77 | 28 | 51.0 |
| MPS 6 | MPS IIIC | M | 33 | 9 | 60.4 |

#### Isolation of GAGs from urine samples

GAGs were extracted from urine samples provided by patients with MPS III and by aged-matched healthy children using biochemical procedures described previously (10,34). Briefly, urine samples were acidified to pH 5.5 with acetic acid and then centrifuged. All the centrifugation steps were performed at 4°C, 2,400 x g for 10 min

The supernatants were incubated with cetylpyridinium chloride (200 µL 5% w/v) for 16h at 4°C and then centrifuged. After successive washes with ethanol solutions saturated with sodium chloride, ethanol, and ether, the pellets were dried and resuspended in 0.6 M NaCl at 4°C for 3h. After centrifugation, supernatants containing solubilized GAGs were precipitated with ethanol for 16h at 4°C. The pellets were washed as described above, resuspended in 0.6 M NaCl, and used as GAG fractions.

Microglial cell line culture and treatment with GAGs or LPS

#### Cell culture

The BV-2 murine microglial cell line was maintained at a density of 0.05×10^6^ cells/ml in BV-2 complete medium (RPMI-1640-Glutamax (RPMI) medium supplemented with 5% heat-inactivated fetal bovine serum (FBS), 2 mM glutamine, 1 U/mL penicillin and 1 µg/mL streptomycin) at 37°C in a 5% CO_2_ atmosphere. The BV-2 cells were used from passages 5 to 15 and were negative for mycoplasma.

#### Treatment with GAGs or LPS

GAG fractions from patients with MPS III and from controls were pooled, as the same effects were seen when cells were exposed to fractions from individual patients (as shown previously) (17). GAGs were used at the optimal concentration of 5 μg/ml, as demonstrated previously (10,17).

To check the proinflammatory BV-2 cell response to GAG treatment, cells were seeded for flow cytometry and RT-qPCR assays at a density of 0.1 ×10^6^ cells/well or 0.15×10^6^ cells/well, respectively, in a 6-well plate for 24h before treatment with 5 μg/ml GAGs, as described previously (10,17). RNA was extracted 24h after treatment, and flow cytometry experiments were performed 48h after treatment.

For EV isolation, EV-depleted FBS medium was obtained by the ultracentrifugation of BV-2 complete medium supplemented with 20% FBS for 18h at 100.000 × g and 4°C in 38.5 mL polyallomer Ultra Clear ultracentrifuge tubes (Beckman Coulter), using an Optima XPN-80 ultracentrifuge and an SW 32 Ti Rotor (Beckman Coulter) at maximum acceleration and brake. Cells were seeded 24h before GAG or LPS (0.1 μg/ml) treatment at a density of 0.25×10^6^ cells/ml in 6 x T175 flasks per condition containing 15 mL of EV-depleted medium (5% EV-depleted FBS). Cells were treated for 24h before conditioned medium collection in fresh EV-depleted medium containing GAGs extracted from urine samples of patients with MPS III (MPS III, 5 μg/ml) or age-matched healthy donors (WT, the same volume as in the MPS condition) or with 0.6 M NaCl (CTL, the same volume as in the MPS condition) and filtered on a 0.4 μm filter.

### Isolation and characterization of EVs from conditioned media

All the relevant data from our experiments were sent to the EV-TRACK knowledgebase (EV-TRACK ID: EV231009) (35).

#### Isolation of EVs

Suspensions were centrifuged at 4°C in 50 mL centrifuge tubes (Falcon) with a 5810R centrifuge and an A-4-81 rotor (Eppendorf) and in 38.5 mL polyallomer Ultra Clear ultracentrifuge tubes (Beckman Coulter) with an Optima XPN-80 ultracentrifuge and an SW 32 Ti Rotor (Beckman Coulter) at maximum acceleration and brake.

Procedures were adapted from references (36–38), according to International Society for Extracellular Vesicles guidelines (39) (see Fig. 1A for a detailed diagram of the experiment scheme for ultracentrifugation). To pellet the cells, conditioned medium was centrifuged at 300 × g for 10 min at 4°C (yielding the 0.3K pellet). Supernatant was centrifuged at 2,000 × g for 20 min at 4°C (yielding the 2K pellet), transferred to ultracentrifugation tubes, and ultracentrifuged first for 30 min at 12,000 × g (yielding the 12k pellet) and then for 70 min at 100,000 × g (yielding the 100k pellet). The 12k and 100k pellets were washed in 35 mL of PBS and recentrifuged at the same speed before being resuspended in the appropriate buffer for the downstream experiment (see below). For neuron treatments, the 12,000 x g ultracentrifugation and washing steps were omitted.

**Fig. 1.**
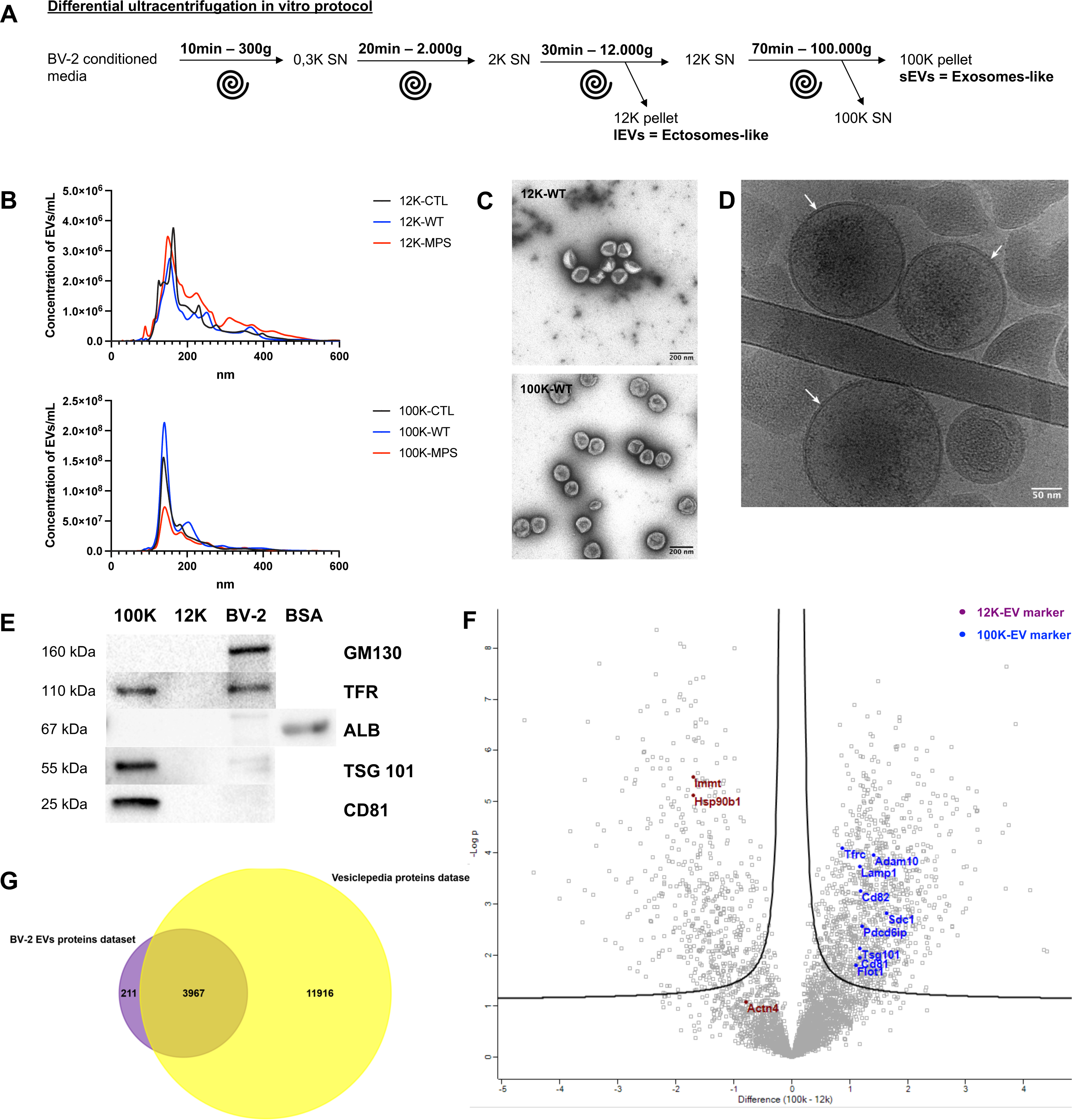
Purification and characterization of EVs from BV-2-cell-conditioned medium. **(A)** Differential centrifugation purification of 12k-EVs and 100k-EVs from BV-2 cell-conditioned medium. **(B)** NTA size distribution graphs of the purified 12k-EVs (upper panel) and 100k-EVs (lower panel) from cell-conditioned media of BV-2 cells treated with WT-GAGs (WT-, n=5), MPS III-GAGs (MPS-, n=5) or vehicle only (CTL-, n=3). The diameter of EVs from the 12k pellet ranges from 100 to 400 nm and that of the EVs from 100k pellet ranges from 120 to 280 nm. The results are expressed as the mean EV concentration per mL for each nm in size. **(C)** Representative images of negative-stained TEM showing pure EV preparations of WT-12k EVs (upper panel) and WT-100k EVs (lower panel). Scale bar = 200 nm. **(D)** Representative cryo-TEM image showing the presence of a lipid bilayer (white arrows) in CTL-100k EVs. Scale bar = 50 nm. **(E)** Western blots performed with CTL-100k EVs (100k), CTL-12k EVs (12k) and BV-2 cell lysates (BV-2) revealed the enrichment of the specific EV protein markers CD81, TSG101, mADAM10, TFR in the 100k pellet and the absence of BSA or GM130 proteins in both pellets. Synthetic BSA (5 μg) as a positive control for the anti-albumin antibody. **(F)** A volcano plot of proteins detected in 100k-EVs vs. 12k-EVs (using a quantitative LC-MS/MS analysis). Mass spectrometry revealed the presence of specific protein markers of large EVs (12k, brown) and small EVs (100k, blue). X axis = the fold-change (100k-12k), y axis = −log10(P value). The p-value was determined in a two-tailed Student’s t-test. Threshold line indicates permutation-based FDR set at 5% combined with fold change set with S0 = 0.1. **(G)** Venn diagram generated using Funrich software, showing that 95% of the identified EVs proteins (in purple) are referenced in the Vesiclepedia public protein database (in yellow). (**C, E – F)** n = 3 biological replicates.

Cells recovered from the 0.3K pellet were pooled with cells detached from the flask by incubation at room temperature in PBS-EDTA (Gibco) and counted in a Malassez cell. Cell viability was assessed in a 0.4% Trypan Blue exclusion assay (Life Technologies). All EV preparations used were cryopreserved at -80°C in sterile (0.22 μm filtered) PBS containing 25 mM trehalose (40,41). The EV concentration per mL was quantified by NTA.

#### NTA

The size distribution and particle concentrations of EV preparations were analyzed using a NanoSight NS300 NTA system (Malvern Panalytical) equipped with a blue laser (488 nm, 70 mW), a CMOS camera (Hamamatsu Photonics), and a syringe pump. EV preparations were resuspended in 100 μL of sterile, filtered PBS containing 25 mM trehalose and stored at -80°C before the NTA experiment. Next, the EV preparations were diluted 1:1000 in filtered PBS for the measurement of 20–100 particles/frame under all conditions, injected at a speed of 30 arbitrary units into the measurement chamber. The EV flow was recorded during triplicate measurements of 60 s each, at 20°C. The data acquisition settings were the same in all experiments, with the camera level set to 9-11, the screen gain set to 7, and the threshold set to 3-4. Data from at least three different experiments were analyzed with NTA 3.2 software (Malvern Panalytical).

#### TEM

EV preparations were resuspended in 100 mM Tris-HCl and stored at 4°C for 24h at most before the TEM experiment. The EV samples were prepared for TEM in a conventional negative staining procedure. Briefly, 10 μL of EV samples were absorbed for 2 min on formvar-carbon-coated copper grids previously ionized using the PELCO easyGlow™ Glow Discharge Cleaning System (Ted Pella Inc., Redding, CA, USA). Preparations were then blotted and negatively stained with 1% uranyl acetate for 1 min. Grids were examined using an 80 kV JEM-1400 electron microscope (JEOL Inc., Peabody, MA, USA), and images were acquired with a digital camera (Gatan Orius, Gatan Inc., Pleasanton, CA, USA).

To analyze the morphology of EVs using cryo-TEM, 3 μL of the EV sample was first deposited onto a glow-discharged 200-mesh lacey carbon grid. Prior to freezing, the grid was loaded into the thermostatic chamber of a Leica EM-GP automatic plunge freezer at 20°C and 95% humidity. Excess solution was blotted from the grid for 1–2 s with Whatman n°1 filter paper, and the grid was immediately flash-frozen in liquid ethane at −185°C. Specimens were then transferred into a Gatan 626 cryo-holder. Cryo-EM was carried out using a Jeol 2100 microscope equipped with a LaB6 cathode operating at 200 kV under low-dose conditions. Images were acquired using SerialEM software (Mastronarde, 2005), with the defocus ranging from 600 to 1000 nm, using a Gatan US4000 CCD camera. This device was placed at the end of a GIF quantum energy filter (Gatan Inc. Berwyn, PA, USA) operating in zero-energy-loss mode, with a slit width of 25 eV. Images were recorded at a magnification corresponding to a calibrated pixel size of 0.87 Å.

### Flow cytometry analysis

After GAG treatment, cells were detached and resuspended in culture supernatant. The total resuspension volume was aliquoted into two tubes (for stained cells and unstained control cells). After the cells had been recovered by centrifugation at 400 × g for 5 min at 4°C, they were washed in PBS with 5% FBS and recentrifuged. Cell surface markers were labelled with fluorophore-labeled primary antibodies at appropriate dilutions (see table 2) in PBS with 5% FBS during 20 min at 4°C. Cells were washed in PBS with 5% FBS, centrifuged at 400 × g for 5 min at 4 °C, and resuspended in 300 µL of PBS before flow cytometry acquisitions. All the conditions were measured on a four-laser custom LSR II-Fortessa flow cytometer (Beckton Dickson). Controls were performed for each experimental sample, including unstained cells and manual compensations during measurements. The acquisition was gated using FlowJo software (BD). The fluorescence was quantified as the mean fluorescence intensity (MFI) ratio between stained and unstained cells. The threshold for statistically significant differences in the MFI ratio was p<0.05 in an unpaired, ordinary, one-way ANOVA of data from at least four different experiments.

**Table 2:**
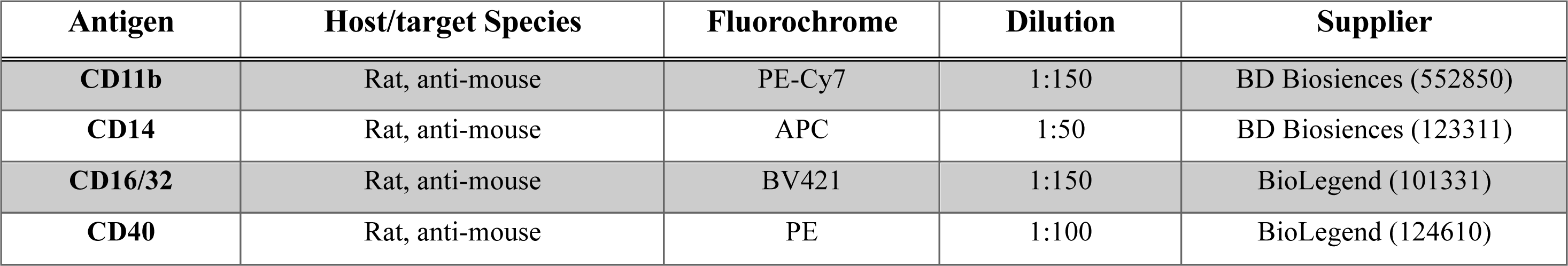
Flow cytometry antibodies.

| Antigen | Host/target Species | Fluorochrome | Dilution | Supplier |
| --- | --- | --- | --- | --- |
| CD11b | Rat, anti-mouse | PE-Cy7 | 1:150 | BD Biosciences (552850) |
| CD14 | Rat, anti-mouse | APC | 1:50 | BD Biosciences (123311) |
| CD16/32 | Rat, anti-mouse | BV421 | 1:150 | BioLegend (101331) |
| CD40 | Rat, anti-mouse | PE | 1:100 | BioLegend (124610) |

### Western blotting and mass spectrometry proteomic analysis

#### Protein extraction

EV preparations and 4 × 10^6^ BV-2 cells were lysed in RIPA lysis buffer 1X (Sigma) supplemented with protease inhibitor cocktail (Roche) for 20 min on ice, centrifuged at 18,500 × g for 15 min, and stored at -20°C before western blotting experiments or at -80°C before mass spectrometry proteomic experiments.

#### Western blotting

The protein concentration in cell lysates was assayed using a BCA gold kit (Life Technologies). Proteins were denatured (or not) in Laemmli buffer under reducing conditions (5% β-mercaptoethanol) and then heated for 5 min at 95°C. 5 µg of proteins from cell lysate or a quarter of the EV protein extract were loaded onto 4–15% Mini-PROTEAN® TGX™ Precast Protein Gels (Biorad) in Tris-glycine buffer for electrophoresis at constant amperage (20 mA per gel) for 1h. Proteins were electrotransferred onto nitrocellulose membranes using the trans-blot turbo transfer system (constant voltage: 25V, maximum amperage: 1 A; 30 min, Biorad) and membranes were blocked with Tris-buffered saline containing 0.1% Tween 20 (TBST), 5% milk, or bovine serum albumin (BSA) for 2h at room temperature. Membranes were then incubated with primary antibodies in TBST containing 5% milk or BSA overnight at 4°C, followed by incubation with secondary antibodies for 1h at room temperature (see Table 3 for antibody dilutions and blocking conditions). Membranes were washed five times in TBST for 5 min after each incubation step and visualized using the Chemidoc XRS+ Imaging System (LI-COR Biosciences). None of the membrane was stripped.

**Table 3:**
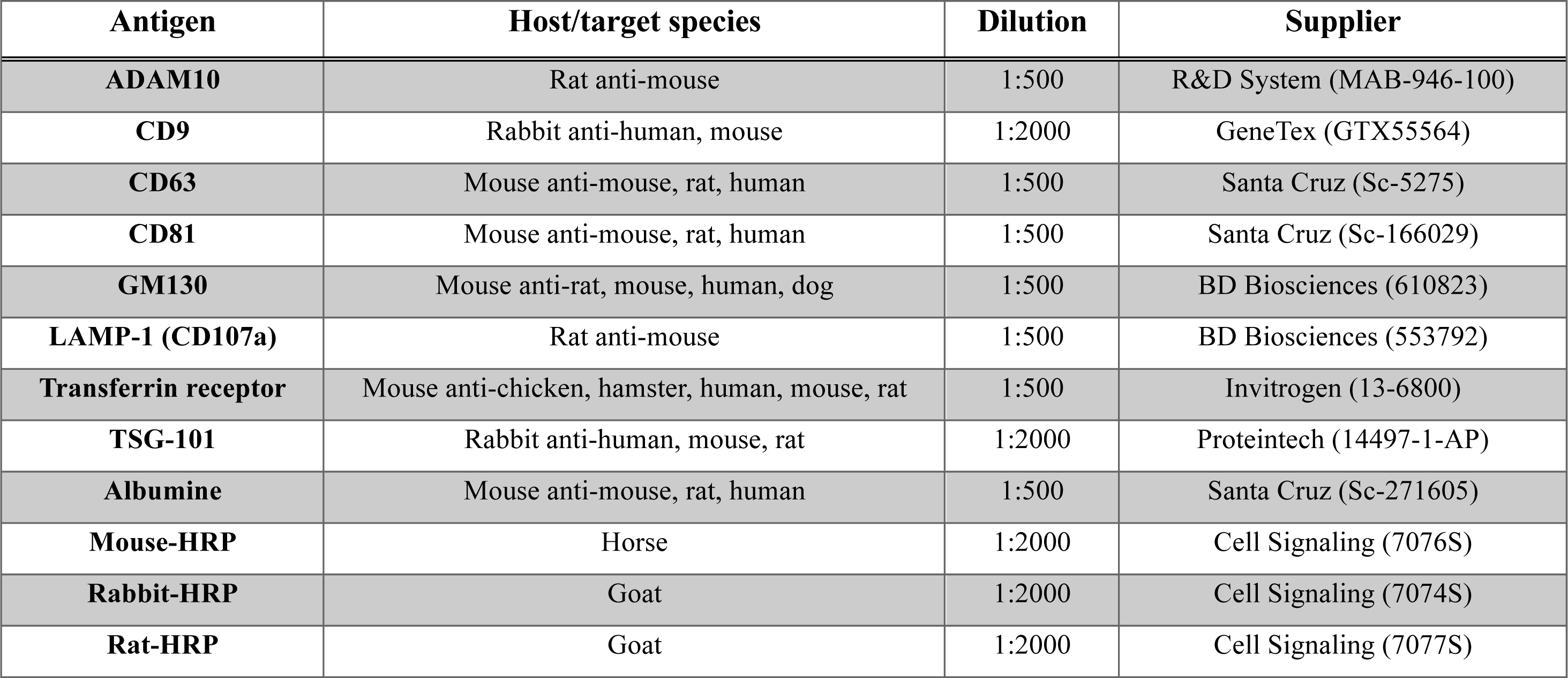
Western blot antibodies.

#### Protein digestion for mass spectrometry

S-Trap^TM^ microspin column (Protifi, Hutington, USA) digestion was performed on 5-10 µg of small or large EV samples, according to the manufacturer’s instructions. Briefly, samples were supplemented with 20% SDS to a final concentration of 5%, reduced with 20 mM Tris(2-carboxyethyl) phosphine hydrochloride and alkylated with 50 mM chloracetamide for 5 min at 95°C. Aqueous phosphoric acid was then added to a final concentration of 2.5%, followed by S-Trap binding buffer (90% aqueous methanol, 100 mM TEAB, pH 7.1). Mixtures were then loaded on S-Trap columns, which were washed five times for thorough SDS elimination. Samples were digested with 1.5 µg of trypsin (Promega) at 47°C for 2h. After elution, peptides were vacuum dried and resuspended in 2% acetonitrile/0.1% formic acid in HPLC-grade water, prior to MS analysis.

#### Nano LC-MS/MS

Tryptic peptides were resuspended in 20 µL, and a volume of 1 µL (for small EVs) or 4 µl (for large EVs) was injected on a nanoelute HPLC system coupled to a timsTOF Pro mass spectrometer (both from Bruker Daltonics, Germany). HPLC separation (solvent A: 0.1% formic acid in water, 2% acetonitrile; solvent B: 0.1% formic acid in acetonitrile) was carried out at 250 nL/min, using a packed emitter column (C18, 25 cm×75 μm 1.6 μm) (Ion Optics, Australia) using a 40 min gradient elution (2 to 11% solvent B for 19 min; 11 to 16% for 7 min; 16% to 25% for 4 min; 25% to 80% for 3 min, and finally 80% for 7 min to wash the column). MS data were acquired using the parallel accumulation serial fragmentation (PASEF) acquisition method in DDA mode. The measurements were carried out over the m/z range from 100 to 1700 Th. The ion mobility value ranged from 0.85 to 1.3 V s/cm^2^ (1/k0). The total cycle time was set to 1.17 s, and the number of PASEF MS/MS scans was set to 10.

#### Data analysis

Data were analyzed using MaxQuant version 2.0.1.0 and searched with Andromeda search engine against the UniProtKB/Swiss-Prot *Mus musculus* dataset (release 02-2021, 17063 entries). To search for parent mass and fragment ions, the mass deviation was set respectively to 10 ppm and 40 ppm for the main search. The minimum peptide length was set to seven amino acids and strict specificity for trypsin cleavage was required, allowing up to two missed cleavage sites. Carbamidomethylation (Cys) was set as a fixed modification, whereas oxidation (Met) and N-term acetylation (Prot N-term) were set as variable modifications. The FDRs at the peptide and protein level were set to 1%. Scores were calculated in MaxQuant, as described previously (42). The reverse and common contaminant hits were removed from MaxQuant output as well as the protein only identified by site. Proteins were quantified according to the MaxQuant label-free algorithm using LFQ intensities, and protein were quantified using at least one peptide per protein. Lastly, matching between runs was allowed.

Statistical and bioinformatic analysis, including hierarchical clustering, heatmap and principal component analysis, were performed with Perseus software (version 1.6.14.0)(43).

For this purpose, three independent replicates in each of the six groups (100K_Ctrl, 100K_MPS, 100K_WT, 12K_Ctrl, 12K_MPS, 12K_WT) were used for statistical comparisons. The protein intensities were transformed in log2, and proteins identified in at least three replicates in at least one group were tested to statistically (volcano plot, FDR=0.05 and S0=0.5) after imputation of the missing values by a Gaussian distribution of random numbers with a standard deviation of 30% relative to the standard deviation of the measured values and 2.5 standard deviation downshift of the mean. Two-tailed pairwise Student’s t-test was performed in Perseus to compare the 100K and the 12K conditions (S0 = 0.1; permutation-based FDR = 5%, 9 independant replicates in each condition). To study the global differences that were present in one of the 6 groups, 1-way ANOVA was performed in Perseus (S0 = 0.1; FDR = 5%, 3 independant replicates in each condition). Then, significant proteins were filtered and log2 LFQ intensities were z-scored before heatmap hierarchical clustering analysis.

### RT-qPCR and small RNA sequencing

#### RNA extraction

For RT-qPCR experiments of mRNAs, total RNAs from BV-2 cells were extracted using TRIzol reagent (Invitrogen), according to the manufacturer’s instructions. The purified RNAs were eluted into 30 μL nuclease-free water, quantified using a Nanoview spectrophotometer with a μCuvette (Eppendorf), and stored at −80°C until analysis.

For RNA sequencing and RTqPCR of miRNAs, total RNAs from EV preparations and 0.5 × 10^6^ cells were extracted using the miRNeasy Tissue/Cells Advanced Micro Kit (QIAGEN), according to the manufacturer’s instructions. The purified RNAs were eluted into 20 μL nuclease-free water, quantified using Nanodrop 2000 spectrophotometer (ThermoFisher Scientific), and stored at −80°C until analysis.

#### RT-qPCRs of mRNA and miRNA

mRNAs were reverse-transcribed using an iScript cDNA synthesis Kit (Biorad), according to the manufacturer’s instructions. RT-qPCRs were performed in a thermocycler for 5 min at 25°C, 20 min at 46°C and then 1 min at 95°C to inactivate the reaction. The qPCR primers for genes of interest (Table 4) were used at a concentration of 1 µM and a 1:1 forward:reverse ratio. Amplification of first-strand cDNAs was performed on the LightCycler 480 Real-Time PCR system (Roche), using Light cycler 480 SYBR Green I Master mix 2X (Roche) in 96-well plates at 95°C for 10 min, 45 cycles of 95°C for 10 s, and 72°C for 30 s, followed by a melting curve analysis at 65–95°C. Gene expression level differences were considered to be statistically significant if they had a p-value <0.05 in an unpaired t-test. Two to five independent replicates were analyzed in technical duplicates.

**Table 4:**
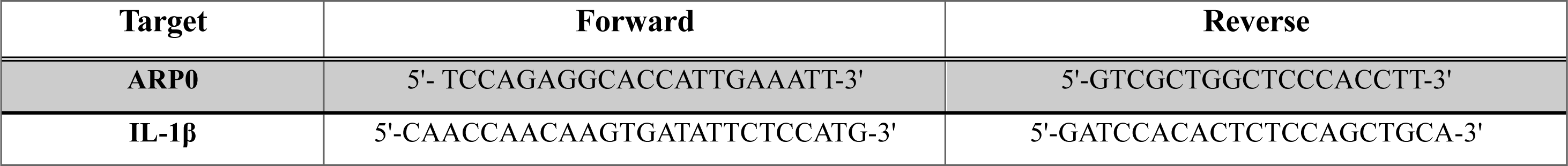
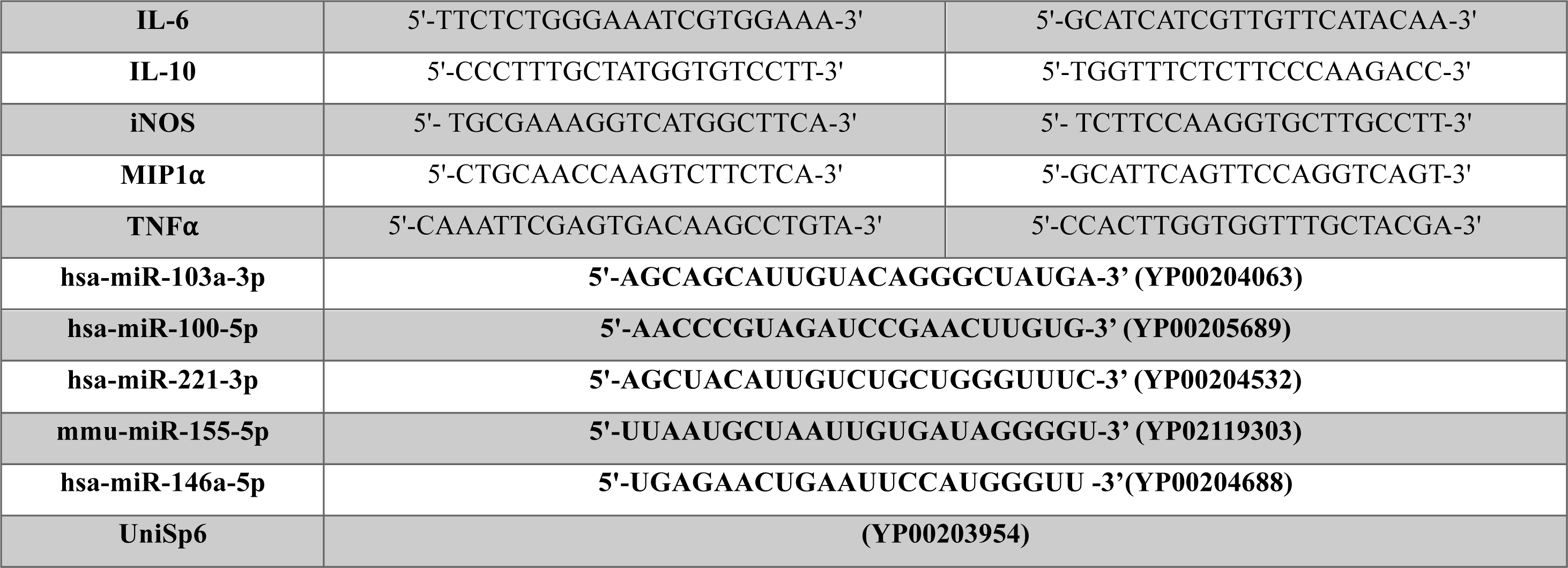
RT-qPCR primers.

For miRNAs, RT and amplification was performed using miRCURY LNA SYBR Green PCR Kit (QIAGEN), according to the manufacturer’s instructions. For RT, a synthetic RT spike-in internal control (UniSp6) was added, and reactions were incubated in a thermocycler for 60 min at 42 °C and then 5 min at 95 °C to inactivate the reaction. Amplification of first-strand cDNAs was performed on a LightCycler 480 Real-Time PCR system (Roche) in 96-well plates at 95 °C for 2 min and 45 cycles of 95 °C for 10 s and 56 °C for 60 s, followed by melting curve analysis at 60–95 °C. The expression levels of the endogenous miRNAs and spike-ins were examined using the miRCURY LNA miRNA PCR Assay (QIAGEN), using primers designed for the *Homo sapiens* genome and validated for the *Mus musculus* genome (Table 4). Gene expression–level differences were considered to be as statistically significant if they had a p-value <0.05 in an ordinary one-way ANOVA. Four independent replicates were analyzed in technical duplicates.

For both qPCR analyses, additional controls were performed for each cDNA amplification: the assessment of amplification efficiency, and the detection of possible primer dimerization via an analysis of dissociation curves. The cycle threshold (Ct) value was determined as the number of PCR cycles at which specific amplification of the target sequence occurred. A Ct >38 was considered to be the background signal. The amount of cDNA was expressed as ΔCt, the difference between the Ct measured for the amplification of the examined cDNA and the reference Ct measured for the amplification of ARP0 or miR-103a-3p cDNAs. The fold change was expressed as ΔΔCt, which is the difference between the ΔCt in the treated condition and the ΔCt of the control condition.

#### Library preparation and RNA sequencing

RNA was quantified using the Qubit RNA HS Assay Kit on the Qubit Fluorometer (Invitrogen). The quality was evaluated on an Agilent 4150 TapeStation with the High Sensibility RNA ScreenTape kit (Agilent). Each sample’s libraries were built with the QIAseq miRNA UDI Library Kit (QIAGEN), according to the manufacturer’s instructions and quantified using the Qubit 1X dsDNA HS kit on the Qubit Fluorometer (Invitrogen). All libraries were sequenced using a single-read strategy (10 million reads per sample) on an NextSeq550 platform, with 150 cycles of the MidOutput flowcell (Illumina), a read length of 72 bp, and dual indices of 10 bp. Raw data were converted to FASTQ files using bcl2fastq (v2.20).

#### Data analysis

Three independent replicates were analyzed using QIAGEN RNA-seq Analysis Portal 4.1 (QIAGEN, Aarhus, Denmark) and the analysis workflow (v1.2) and using the Legacy Analysis Pipeline on the QIAGEN GeneGlobe portal. The sequences were aligned against the reference miRBase_v22 and the *Mus musculus* genome (GRCm38.101). Gene expression–level differences were accepted as statistically significant when the p-value in an ANOVA was <0.05 and when the fold change was between 1 and -1 after an intergroup comparison.

### Bioinformatics analyses

#### PPIs and GO annotation

Cytoscape software (v3.10.1) and the STRING public database (v12.0) were used to generate local proteins networks and map protein data onto a PPI network. No more than 10 predicted functional interactors were added in the first shell, using a medium level of confidence (0.4) for edge prediction. For proteins without interactions, pertinent GO biological processes were attributed manually when FDR <0.05 and strength >1.

#### miRNA target filters and canonical pathway analysis

IPA (Ingenuity Pathway Analysis, Winter Release December 2023, QIAGEN, Aarhus, Denmark) was used to perform and miRNA Target Filter analysis, according to the software’s manual. The analysis considered all relationship sources in the mouse and the human, and the following filters were applied: miRNA confidence (Experimentally observed, High (predicted)), Tissue/Cell Line (Astrocytes, Macrophages, Neurons, Nervous system, CNS cell lines, Neuroblastoma cell lines), Pathway (Neurotransmitters and other nervous system signaling), Disease (Neurological disease, Inflammatory response, inflammatory disease, developmental disorder). The miRNA Target Filter analysis with a published dataset (21) was performeds as follow: miRNA confidence (Experimentally observed, High (predicted)), Expression pairing (miRNA upregulated, mRNA downregulated). A standard IPA core analysis (including canonical pathways) was performed with Fisher’s exact test (p<0.05) and application (or not) of the “Neurotransmitters and other nervous system signaling” filter. Networks involving miRNAs and mRNAs were designed with the Path Designer tool from IPA (Winter Release December 2023, QIAGEN, Aarhus, Denmark).

#### Culture and treatment of primary cortical neurons

C57BL/6J mice were purchased from Charles River Laboratories (Saffron Walden, UK). Housing, care, coupling and gestation were performed in the US 006/CREFRE INSERM/UPS - Service Zootechnie Expérimentation Purpan animal facility (Toulouse, France), in accordance with French regulations and the European Union Council Directive (2010/63/EU). Mouse experiments were performed by authorized investigators and were approved by the government-accredited animal care and use committee (*Comité d’Ethique de l’US 006/CREFRE* (Toulouse, France); reference number: PI-U1291-JA-37).

Primary cortical neurons were prepared from newborn C57BL/6J mice, using a protocol adapted from (44) The neocortices were isolated from pups on day 0-1 in cold Earle’s Balanced Salt Solution (EBSS) and incubated for 15 min at 37°C in EBSS containing 10 U/ml papain (Worthington), followed by gentle dissociation in PBS containing 1.5 mg/ml BSA, 1.5 mg/ml trypsin inhibitor (Sigma-Aldrich), and 66 µg/ml DNase I (Life Technologies). The cells were filtered twice through a 4% BSA cushion by centrifugation, and seeded at a density of 350000 cells/well in 12-well plates with 12 mm diameter glass coverslips. The plates and coverslips had been coated with 1 mg/mL poly-D-lysine (Merck) and 4 μg/mL laminin (Life Technologies).

#### Treatment of neurons with microglial EVs

Primary cortical neurons were maintained for 7 days (DIV 7) before EV treatment at 37°C in 5% CO_2_ and cultured in serum-free Neurobasal medium (Life Technologies) supplemented with 1.2% GlutaMAX-I (Life Technologies), 1.2 U/mL of penicillin, 1.2 µg/mL streptomycin, and 2% B-27 supplement (Life Technologies). Neurons were treated from DIV 7 to DIV 9 with 8.75 x 10^8^ EVs (WT-, MPS- or LPS-EVs) per well or with vehicle every 24h for 72h.

Coverslips (in duplicate for each condition) were fixed after 24h (DIV 8), 48h (DIV 9) or 72h (DIV 10) of treatment in 4% paraformaldehyde and 4% sucrose solution in PBS for 20 min at room temperature and stored at 4°C in PBS prior to immunostaining.

### Immunocytochemistry

#### Immunostaining

After fixation, cells were permeabilized with 0.1% Triton 100X in PBS 1X for 10 min at room temperature and washed once with PBS 1X. Neurons were blocked with 3% donkey serum and 3% Bovine Serum Albumine (BSA) in PBS 1X for 1h at room temperature and incubated overnight at 4°C with primary antibodies. On the next day, coverslips were washed three times in PBS 1X before staining with secondary antibodies for 1 h at room temperature (see Table 5 for details of antibodies and their dilutions). The terminal deoxynucleotidyl transferase dUTP nick-end labeling (TUNEL) assay was performed according to the manufacturer’s recommendations. Lastly, nuclei were washed three times in PBS 1X for 5 min at room temperature, stained with Hoechst 33342 (Life Technologies) diluted 1:2000 in PBS 1X, and washed again three times in PBS 1X for 5 min. Coverslips were mounted on slides with a few microliters of ProLong™ Gold Antifade Mountant (Life Technologies) and dried either overnight at room temperature or longer at 4°C.

**Table 5:** Immunocytochemistry antibodies.

| Antigen | Host/target species | Dilution | Supplier |
| --- | --- | --- | --- |
| **MAP2** | Chicken anti-mouse, rat | 1:500 | Abcam (ab5392) |
| **β-III Tubulin** | Mouse anti-human, mouse, rat, chicken | 1:100 | R&D System (MAB1195) |
| **Chicken-Alexa Fluor<sup>TM</sup> 568** | Goat | 1:1000 | Invitrogen (A11041) |
| **Mouse-Alexa Fluor<sup>TM</sup> 647** | Donkey | 1:1000 | Invitrogen (A32787) |

#### Image analysis

All the slides were imaged using a Zeiss ApoTome 2 (Carl Zeiss Group, Oberkochen, Germany) fluorescence microscope. Images of neurites and TUNEL assay results were analyzed as 25-square stitched mosaics with a 10X wide field objective and AI Filament Tracer plug-in (for neurite quantification) or a colocalization module (for TUNEL quantification) in Imaris software (version 10.0, Oxford Intruments, Abingdon-on-Thames, Royaume-Uni). Neurite variables were expressed as differences between WT or MPS III conditions and CTL condition. Images for dendrite morphology analysis were obtained with 40X and 63X oil immersion objectives, processed using Zeiss ZEN software, and quantified using ImageJ2 2.3.0. The results of at least four different experiments (with different neurons and EV batches) were quantification on blind basis, using the same parameters for each experiment.

The dendritic arborization was quantified in a Sholl analysis with an “inside-out” scheme (45).

### Statistical analyses

All statistical tests were performed with GraphPad Prism software (version 10.1.0, GraphPad Software LLC, San Diego, CA, USA). Intergroup differences were analyzed using an independent-sample T-test. Three or more groups were compared in a one-way ANOVA with Šidák correction or in a two-way ANOVA. The threshold for statistical significance was set to p<0.05, unless otherwise stated. Data were expressed as the mean ± SEM.

## Results

### Pro-inflammatory activation of BV-2 cells after treatment with pathogenic GAGs extracted from the urine of patients with MPS III

In order to obtain a sufficient amount of a homogenous population of microglia derived-EVs for the comprehensive characterization of protein and small RNA contents and for a primary cortical neuron uptake assay, we used an *in vitro* model of MPS III: the BV-2 mouse microglial cell line treated with pathogenic GAGs extracted from the urine of patients with MPS III (henceforth referred to as MPS-GAGs). We first checked that BV-2 cells could be activated by 5 µg/mL MPS-GAGs, as used previously with primary microglia and neurons (10,17). Using quantitative reverse transcription PCR (RT-qPCR, Additional file 1: Fig. S1A) and flow cytometry (Additional file 1: Fig. S1B), we showed that BV-2 cells expressed more pro-inflammatory cytokines (MIP1⍺, IL-6, IL-1β, and TNF⍺) and surface markers (CD14, CD40, CD16/32 or CD11b) after exposure to MPS-GAGs.

We next purified and characterized the large and small EVs secreted by BV-2 cells after treatment with either MPS-GAGs, urinary GAGs from healthy children (WT-GAGs), or GAG resuspension buffer (the control (CTL) condition).

### Characterization of EVs

As specified in the guidelines from the International Society for Extracellular Vesicles (39,46), the vesicles in pellets obtained after centrifugation at 12,000 x g (henceforth the 12k pellet enriched in large, ectosome-like EVs with an expected diameter > 150 nm) and 100.000 x g (the 100k pellet enriched in small, exosome-like EVs with an expected diameter of 30-150 nm) (Fig. 1A) were analyzed with regard to particle size and concentration, using nanoparticle tracking analysis (NTA) and transmission electron microscopy (TEM) (Fig. 1B, C and D). The presence of EV protein markers was checked using Western blotting and mass spectrometry (Fig. 1E and F).

#### Nanoparticle tracking analysis

In a comparison of CTL, WT-GAG and MPS-GAG EVs, the NTA showed that when compared to the 12k pellets, all the 100k pellets had a higher total particle concentration. As expected, the particle size was smaller (for both the average particle size (mean) and the peak particle size (mode)), indicating that they are enriched in small EVs (Table 1 and Fig. 1B). On the other hand, 12k pellets showed a higher and more heterogenous particle size (for both mean and mode particle size) when compared to the 100k pellets, suggesting that 12k pellets are depleted in small EVs (Table 1 and Fig. 1B). Also, when considering GAGs treatments, we did not notice any difference in size or concentration between the conditions.

**Table 1.** Results of the NTA.

| Conditions |  | Mode (nm) | Mean (nm) | Total concentration (EVs/mL) |
| --- | --- | --- | --- | --- |
| CTL | 12K | 164 ± 13 | 210 ± 14 | 3E+08 ± 1E+08 |
| | 100K | 139 ± 3 | 180 ± 6 | 7E+09 ± 6E+08 |
| WT | 12K | 151 ± 6 | 216 ± 23 | 2E+08 ± 2E+08 |
| | 100K | 140 ± 3 | 182 ± 8 | 1E+10 ± 8E+09 |
| MPS | 12K | 147 ± 7 | 234 ± 28 | 4E+08 ± 2E+08 |
| | 100K | 141 ± 5 | 197 ± 5 | 5E+09 ± 2E+09 |

#### Transmission electron microscopy

The purified EVs were examined by TEM. Negative staining revealed pure EVs preparation in both 100k and 12k pellets (Fig. 1C). The particle size and quantity were consistent with that determined in the NTA. Cryo-TEM confirmed the presence of a membrane bilayer - a hallmark of EVs (Fig. 1D).

#### EV protein marker analysis, using Western blotting and mass spectrometry

We used Western blotting of the BV-2 control condition, whole cell lysate, and 12k and 100k pellets to establish the presence of typical transmembrane or lipid-bound protein markers of EVs (CD9, CD63, CD81, LAMP1, TfR) and cytoplasmic protein markers (ADAM10 and Tsg101) (Fig. 1E and Additional file 1: Fig. S2A) (39,46). Tetraspanin, CD81, TfR and TSG 101 were highly enriched in the 100k pellet (Fig. 1E). To assess the degree of purity of our EV preparations, we tested for albumin because the latter is supposedly the best negative marker for EVs isolated from cells cultured in the presence of bovine serum (39). Albumin was only detected in BV-2 cell lysate; this observation suggested that our 100k EV preparation was highly pure and was consistent with the TEM images (Fig. 1C). Moreover, the Golgi apparatus marker GM130 was found in the cell lysate fraction only (Fig. 1E).

Due to the small number of particles in the 12k pellet, we were able to evaluate protein markers of large EVs in an LC-MS/MS analysis only. A volcano plot of the overall distribution of 12k and 100k proteins showed that many proteins were significantly enriched (according to the criteria (log2 |fold-change| ≥ 1.2 and *p* < 0.05)) in each subpopulation of EVs (Fig. 1F). The differentially expressed proteins included specific markers of small EVs and specific markers of large EVs. The EV protein markers identified in an LC-MS/MS analysis are listed in Table S1 (Additional file 1). A principal component analysis discriminated well between different EV sources (12k vs. 100k) (Additional file 1: Fig. S2B) and confirmed that the EV isolation and label-free quantitative proteomics protocols were highly reproducible. A Venn diagram showed that 3967 (95%) of the 4178 identified proteins were present in the Vesiclepedia public database (Fig. 1G).

Given that the characterization of the EVs from 12k and 100k pellets gave satisfactory results, we pursued the experiments by evaluating the effect of WT-GAGs and MPS-GAGs on EVs’ proteome and small RNA content.

### Differentially abundant proteins in microglia derived-EVs treated with MPS-GAGs

The 12k and 100k pellets obtained after BV-2 cell treatment with WT-GAGs, MPS-GAGs or CTL buffer (i.e. six conditions in total) were analyzed using label-free, quantitative proteomics. We identified a total of 4332 unique proteins. The numbers of proteins were similar in all six conditions. We next sought to identify the significantly altered EVs proteins among the six conditions by applying a one-way ANOVA (S0 = 0.1 permutation-based FDR < 0.05). In total, 43 significantly altered proteins were identified in an unsupervised hierarchical clustering analysis of the differentially expressed proteins. We observed that only 13 proteins were more abundant in the 12k pellet than in the 100k pellet. Interestingly, the biological replicates from both 12k and 100k MPS-EVs were grouped together in one cluster, while the CTL-EVs and WT-EVs replicates were grouped together in another cluster; this suggested that WT-GAGs do not modify the EVs’ protein content (Fig. 2A). However, relative to WT-EVs or CTL-EVs, 11 proteins were significantly upregulated and 19 proteins were significantly downregulated in both 12k and 100k MPS-EVs (Table 2).

**Fig. 2.**
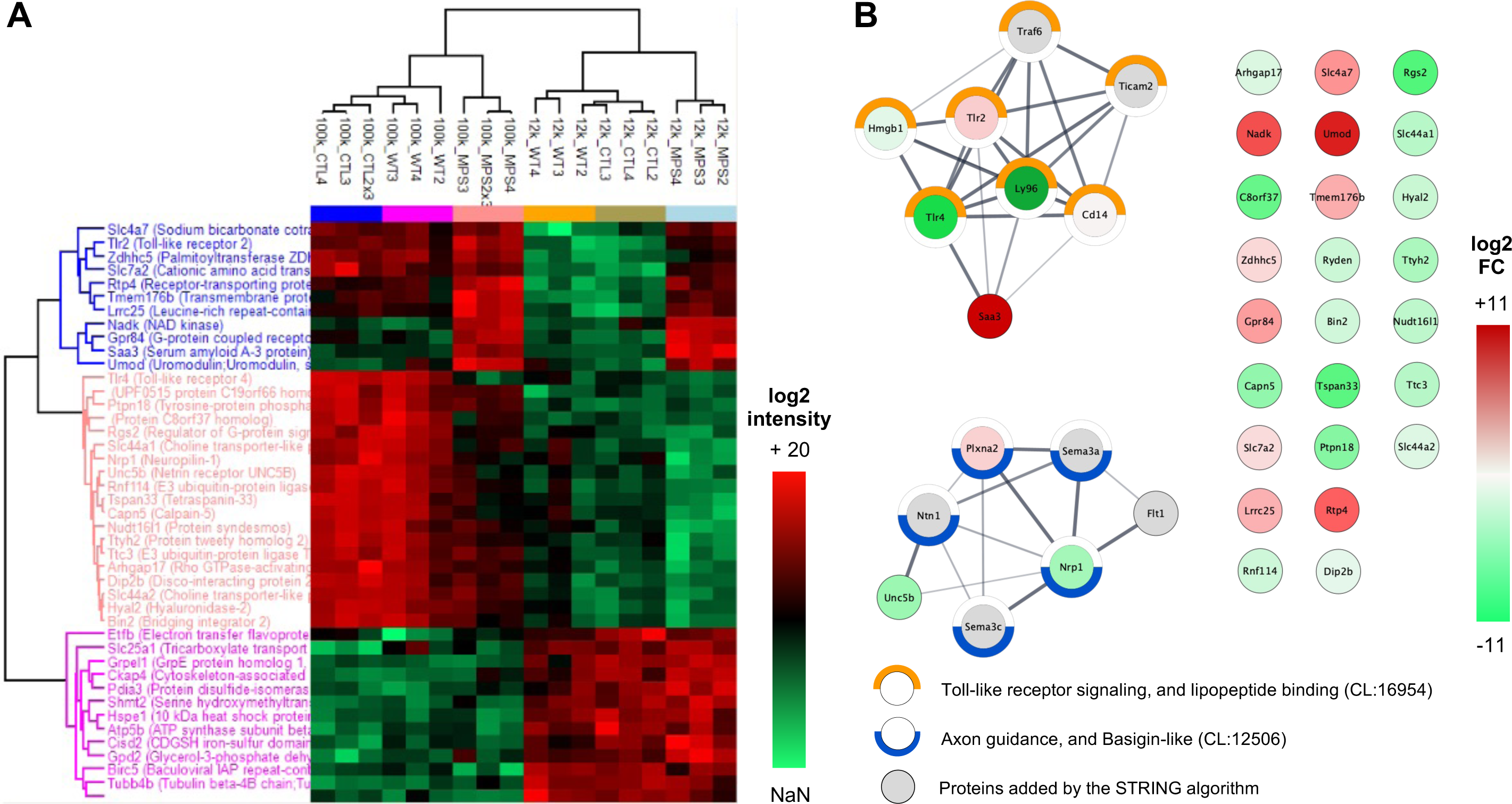
Quantitative proteomics: a comparison of EVs released by microglia treated with WT vs. MPS III-GAGs. **(A)** A heat map of a quantitative differential analysis (LC-MS/MS) of proteins present in EVs as a function of the GAG treatment and type of pellet. EVs were collected from conditioned media of BV-2 cells treated with WT-GAGs (_WT), MPS III-GAGs (_MPS) or vehicle only (_CTL) and proteins extracted from the 12k pellet (12k_) or the 100k pellet (100k_). Log2 fold-change, using label-free quantified intensity. The q-value was determined in a one-way ANOVA performed with Perseus software. Heat colors were scaled per marker (high expression in red; low expression in green). n=3 biological replicates. **(B)** Protein-protein interaction (PPI) networks of the 30 proteins differentially expressed when comparing MPS III-EVs with WT-EVs; the proteins were part of the *Toll-like receptor signaling, and lipopeptide binding* and the *Axon guidance, and basigin-like* networks. The networks were obtained using the STRING database and Cytoscape software. PPI enrichment p-value = 7.41 x10^-6^. The color of the bubble represents the mean log2(fold-change) of proteins expressed in MPS EVs compared with pooled CTL EVs and WT EVs (high expression in red; low expression in green). Half circles around bubbles correspond to STRING local network clusters, *Toll-like receptor signaling, and lipopeptide binding* (in orange, FDR: 3.52x10^-10^) and *Axon guidance, and basigin-like* (in blue, FDR: 2.5x10^-4^).

**Table 2.** Gene ontology annotation of proteins differentially expressed when comparing MPS-EVs with WT-EVs.

| Gene names | log2 FC | -log10 FDR | Uniprot IDs | Protein names | Gene ontology biological process |
| --- | --- | --- | --- | --- | --- |
| **Saa3** | 10,979 | 5,000 | P04918 | Serum amyloid A-3 protein | Inflammatory response; GO:0006954, Chemotaxis; GO:0006935, Positive regulation of reactive oxygen species metabolic process, GO:2000379; Positive regulation of neuron death; GO:1901216 |
| **Umod** | 8,719 | 1,263 | Q91X17 | Uromodulin;Uromodulin, secreted form | Apoptotic process; GO:0006915, Response to oxygen-containing compound; GO:1901700, Cellular response to tumor necrosis factor; GO:0071356 |
| **Nadk** | 5,190 | 0,963 | P58058 | NAD kinase | Negative regulation of apoptotic process; GO:0043066, NADP metabolic process; GO:0006739 |
| **Rtp4** | 4,013 | 1,248 | Q9ER80 | Receptor-transporting protein 4 | Innate immune response; GO:0045087, Positive regulation of cytokine production; GO:0001819 |
| **Slc4a7** | 2,426 | 1,307 | Q8BTY2 | Sodium bicarbonate cotransporter 3 | Regulation of intracellular pH; GO:0051453 |
| **Gpr84** | 2,162 | 1,164 | Q8CIM5 | G-protein coupled receptor 84 | Regulation of cytokine production; GO:0001817, Response to oxygen-containing compound; GO:1901700, Inflammatory response; GO:0006954 |
| **Tmem176b** | 1,780 | 1,270 | Q9R1Q6 | Transmembrane protein 176B | Negative regulation of dendritic cell differentiation; GO:2001199 |
| **Lrrc25** | 1,695 | 1,233 | Q8K1T1 | Leucine-rich repeat-containing protein 25 | Cell-cell adhesion via plasma-membrane adhesion molecules; GO:0098742, Regulation of cytokine production; GO:0001817 |
| **Tlr2** | 1,022 | 1,212 | Q9QUN7 | Toll-like receptor 2 | Positive regulation of cytokine production; GO:0001819, Inflammatory response; GO:0006954, Cellular response to oxygen-containing compound; GO:1901701 |
| **Zdhhc5** | 0,749 | 1,317 | Q8VDZ4 | Palmitoyltransferase ZDHHC5 | Protein acylation; GO:0043543, Protein palmitoylation; GO:0018345, Regulation of postsynaptic membrane neurotransmitter receptor levels; GO:0099072 |
| **Slc7a2** | 0,681 | 1,175 | P18581 | Cationic amino acid transporter 2 | Carboxylic acid transmembrane transport; GO:1905039, Innate immune response; GO:0045087, Regulation of leukocyte cell-cell adhesion; GO:1903037 |
| **Dip2b** | -0,287 | 1,285 | Q3UH60 | Disco-interacting protein 2 homolog B | Mitochondrial respiratory chain complex I assembly; GO:0032981, mRNA transport; GO:0051028 |
| **Slc44a2** | -0,408 | 1,266 | Q8BY89 | Choline transporter-like protein 2 | Choline transmembrane transporter activity; GO:0015220, Cell adhesion; GO:0007155 |
| **Arhgap17** | -0,452 | 1,281 | Q3UIA2 | Rho GTPase-activating protein 17 | Actin cytoskeleton organization; GO:0030036, Establishment or maintenance of cell polarity; GO:0007163, Cell migration; GO:0016477, Vesicle-mediated transport; GO:0016192, Dendritic spine morphogenesis; GO:0060997 |
| **Bin2** | -0,636 | 1,298 | D3Z6Q9 | Bridging integrator 2 | Vesicle-mediated transport; GO:0016192, Synaptic vesicle endocytosis; GO:0048488, Dendritic spine morphogenesis; GO:0060997 |
| **Rnf114** | -0,675 | 1,273 | Q9ET26 | E3 ubiquitin-protein ligase RNF114 | Protein ubiquitination; GO:0016567, Apoptotic process; GO:0006915, Regulation of I-kappaB kinase/NF-kappaB signaling; GO:0043122 |
| **Ryden** | -0,710 | 1,278 | Q8CAK3 | UPF0515 protein C19orf66 homolog |  |
| **Hyal2** | -0,744 | 1,291 | O35632 | Hyaluronidase-2 | Glycosaminoglycan catabolic process; GO:0006027, Regulation of cell migration; GO:0030334, Negative regulation of apoptotic signaling pathway; GO:2001234 |
| **Nudt16l1** | -0,918 | 1,246 | Q8VHN8 | Protein syndesmos | Double-strand break repair; GO:0006302, Mitochondrial transport; GO:0006839 |
| **Slc44a1** | -1,019 | 1,264 | Q6X893 | Choline transporter-like protein 1 | Nitrogen compound transport; GO:0071705, Early endosome to late endosome transport; GO:0045022, Endosome to lysosome transport; GO:0008333 |
| **Ttc3** | -1,035 | 1,328 | O88196 | E3 ubiquitin-protein ligase TTC3 | Wnt signaling pathway; GO:0016055 |
| **Ttyh2** | -1,233 | 1,175 | Q3TH73 | Protein tweety homolog 2 | Intra-Golgi vesicle-mediated transport; GO:0006891, Cellular response to osmotic stress; GO:0071470, Protein localization to axon; GO:0099612 |
| **Nrp1** | -1,339 | 1,299 | P97333 | Neuropilin-1 | Semaphorin-plexin signaling pathway; GO:0071526, Axonogenesis; GO:0007409, Axon guidance; GO:0007411, Basal dendrite arborization; GO:0150020 |
| **Unc5b** | -1,487 | 1,243 | Q8K1S3 | Netrin receptor UNC5B | Axonogenesis; GO:0007409, Axon guidance; GO:0007411, Anterior/posterior axon guidance; GO:0033564 |
| **Capn5** | -1,623 | 1,129 | O08688 | Calpain-5 | Proteolysis; GO:0006508, Actin cytoskeleton organization; GO:0030036, Response to reactive oxygen species; GO:0000302 |
| **Ptpn18** | -1,806 | 0,862 | Q61152 | Tyrosine-protein phosphatase non-receptor type 18 | Positive regulation of MAP kinase activity; GO:0043406, ERBB2-EGFR signaling pathway; GO:0038134, Gliogenesis; GO:0042063 |
| **C8orf37** | -2,337 | 1,066 | Q3UJP5 | Protein C8orf37 homolog |  |
| **Tspan33** | -2,523 | 1,305 | Q8R3S2 | Tetraspanin-33 | Regulation of leukocyte chemotaxis; GO:0002688, Adherens junction organization; GO:0034332, Negative regulation of neuron death; GO:1901215, Complement activation, classical pathway; GO:0006958, Positive regulation of blood-brain barrier permeability; GO:1905605 |
| **Rgs2** | -2,670 | 1,135 | O08849 | Regulator of G-protein signaling 2 | Dopamine receptor signaling pathway; GO:0007212 |
| **Tlr4** | -4,860 | 1,007 | Q9QUK6 | Toll-like receptor 4 | Positive regulation of cytokine production; GO:0001819, Inflammatory response; GO:0006954, Cellular response to oxygen-containing compound; GO:1901701 |

The 30 proteins are ranked by fold change.

#### Protein networks and functional enrichment analysis

Proteins regulate biological processes through functional or physical interactions with other proteins. In a protein-protein interaction (PPI) network analysis, we probed for potential interactions between the 30 differentially expressed proteins and found two PPI networks (Fig. 2B). Functional enrichment analysis of these networks (using the Search Tool for the Retrieval of Interacting Genes/Proteins (STRING)) revealed two clusters: *Toll-like receptor signaling, and lipopeptide binding* (CL:16954, false discovery rate (FDR): 3.5 x 10^-10^) and *Axon guidance, and basigin-like* (CL :12506, FDR : 2.5 x 10^-4^). Since the label-free quantitative proteomic analysis revealed a small number of differentially expressed proteins, the biological processes (Gene Ontology (GO) annotations) involving proteins that were not part of the two networks are reported in Table 2. Most of the proteins that were more abundant in MPS-EVs are involved in the inflammatory response, apoptotic and oxidative processes, macrophage metabolism, or immune cell recruitment. Conversely, most of the proteins that were less abundant in MPS-EVs are involved in neurodevelopment (particularly axonogenesis, synaptogenesis, neuron metabolism, and blood-brain barrier permeability) and, to a lesser extent, the inflammatory response and apoptosis.

### Differences in the abundance of miRNAs in microglia-derived-EVs treated with MPS-GAGs

Total RNA was extracted from the 12k and 100k pellets obtained after treatment of the BV-2 cells with WT-GAGs or MPS-GAGs. The total RNA concentration ranged from 1.01 to 21.2 ng/µl (Additional file 1: Fig. S3A). The mean ± standard error of the mean (SEM) amount of RNA was lower in the 12k pellet than in the 100k pellet (4.05±2.57 vs. 10.2±6.21, respectively). Electrophoresis of the extracted RNA showed that the samples had a high proportion of small RNAs (<200 nt in length) and a low proportion of 18S and 28S rRNAs (Additional file 1: Fig. S3B). The total reads obtained from RNA sequencing and the mappable reads used for alignment were similar in biological replicates from the WT-100k, MPS-100k, WT-12k and MPS-12k pellets (Additional file 2: Table S2). The EVs contained high amounts of various types of small RNAs (mainly tRNAs and miRNAs, and other small non-coding RNAs) and very low amounts of rRNAs and mRNAs. The small RNA distribution pattern was similar in the WT-100k, MPS-100k, WT-12k and MPS-12k pellets (Additional file 1: Fig. S4).

RNA sequencing identified a total of 1960 miRNAs. An ANOVA revealed that levels of 38 miRNAs were significantly altered (i.e. a log2 |fold-change| ≥ 1 and *p* < 0.05) when comparing the four conditions (Table 3 and Fig. 3A). The replicates from the 12k and 100k MPS-EVs were grouped together in one cluster, and the replicates from the 12k and 100k WT-EVs replicates were grouped together in a second cluster. Using unsupervised hierarchical clustering of these 38 differentially abundant miRNAs, we observed that a group of 20 miRNAs was upregulated and a group of 18 miRNAs was downregulated in MPS-EVs, relative to WT-EVs. When we applied more stringent criteria (log2 |fold-change| ≥ 1 and adjusted p < 0.1), a volcano plot of the overall miRNA distribution revealed that MPS-EVs were significantly enriched in four miRNAs (miR-155-5p, miR-146a-5p, miR-221-3p, and miR-100-5p) (Fig. 3B). With regard to the normalized counts per million (CPM) of the four miRNA outliers, miR-100-5p was less abundant (<100 CPM) than the three others (Additional file 1: Fig. S5). The level of expression of these four miRNAs in total cellular RNA and in EV RNA was confirmed by qPCR experiments (Fig. 3C). We found that the most abundant miRNAs in MPS-EVs were also more abundant in extracts from BV-2 cells treated with MPS-GAGs.

**Fig. 3.**
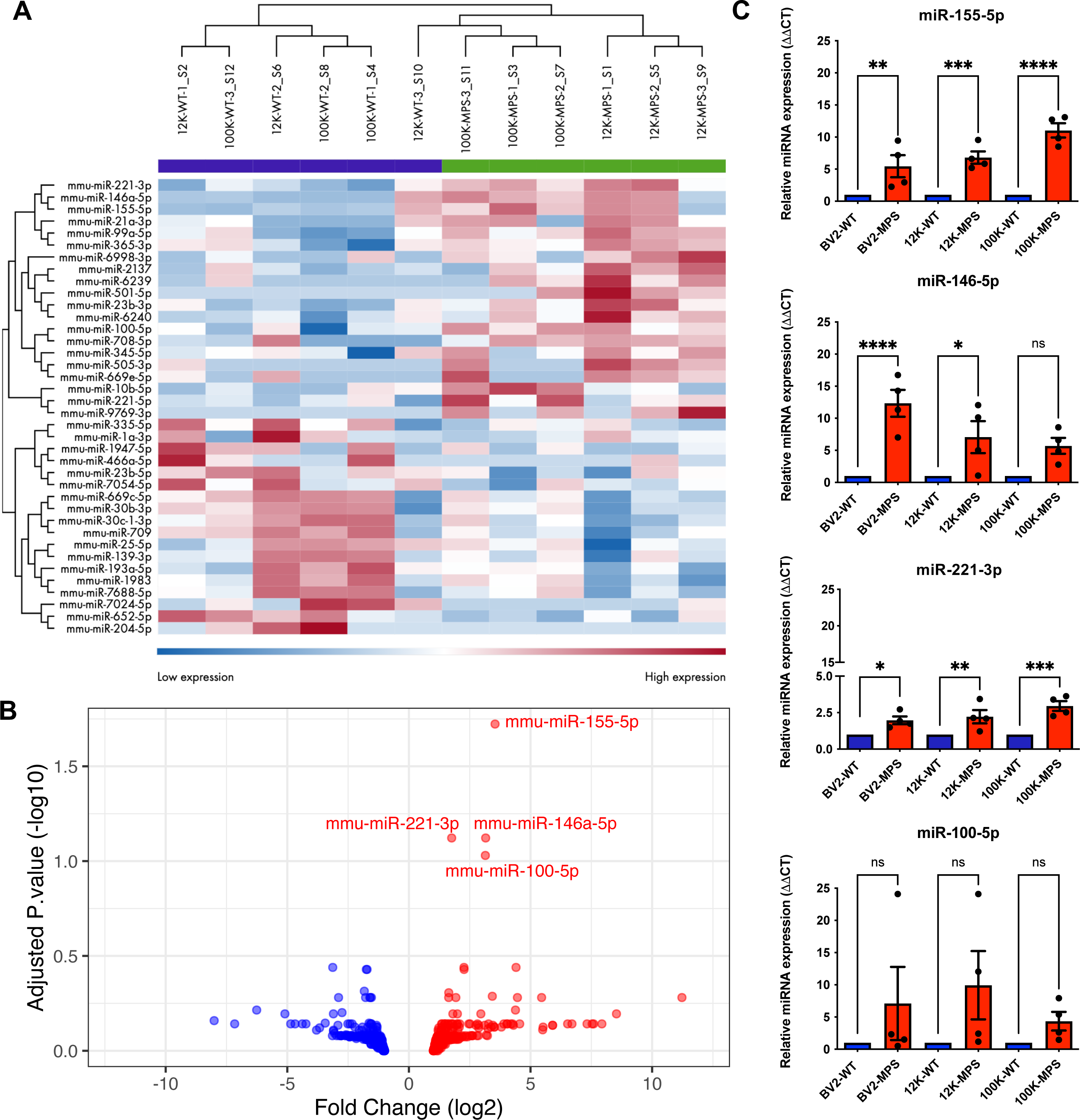
Transcriptomic comparison of miRNAs from EVs released by microglia treated with WT-GAGs vs. MPS III-GAGs. **(A)** A heat map generated in a quantitative differential sequence analysis (Illumina QIAseq) of miRNAs present in EVs, as a function of the GAG treatment and type of pellet. EVs were collected from media conditioned by BV-2 cells treated with WT-GAGs (-WT) or MPS-GAGs (-MPS), and the RNA was extracted from the 12k pellet (12k-) or the 100k pellet (100k-). Log2 fold-change, using RNA sequencing quantified and normalized CPM. The p-value was determined in an ANOVA performed on the QIAGEN RNA-seq Analysis Portal system. The heat colors were scaled for each marker (high expression in red; low expression in blue). n=3 biological replicates. **(B)** A volcano plot of miRNAs detected in MPS III-EV vs. WT-EVs revealed the enrichment of miR-155-5p, miR-146-5p, miR-221-3p and miR-100-5p in MPS-EVs. X axis = log2 |fold-change| ≥ 1 (MPS-WT) with upregulated miRNAs (right, in red) and downregulated miRNAs (left, in blue) in MPS III-EV. y axis = −log10(P value). The p-value was determined in an ANOVA. **(C)** RT-qPCRs of miR-155-5p, miR-146-5p, miR-221-3p and miR-100-5p, represented by bar plots. RNA was extracted from cells (BV-2-) or from EVs (the 12k pellet (12k-) or the 100k pellet (100k-)) collected from conditioned media of BV-2 cells treated with WT-GAGs (-WT, blue) or MPS-GAGs (-MPS, red). One-way ANOVA, n=4 biological replicates (each point represents 1 n), ns = p > 0.05; * = p ≤ 0.05; ** = p ≤ 0.01; *** = p ≤ 0.001; **** = p ≤ 0.0001.

**Table 3.** List of miRNAs expressed differentially when comparing MPS-EVs and WT-EVs.

| miRNA names | log2 FC | -log10 p-value |
| --- | --- | --- |
| mmu-miR-155-5p | 3,54159 | 4,495257364 |
| mmu-miR-146a-5p | 3,15531 | 3,417255034 |
| mmu-miR-221-3p | 1,758177 | 3,585695312 |
| mmu-miR-100-5p | 3,142551 | 3,200246034 |
| mmu-miR-10b-5p | 4,403757 | 2,366632555 |
| mmu-miR-2137 | 2,264235 | 2,374378919 |
| mmu-miR-652-5p | -3,130779 | 2,431211788 |
| mmu-miR-30b-3p | -1,751663 | 2,225556031 |
| mmu-miR-6240 | 2,267543 | 2,201004266 |
| mmu-miR-709 | -1,71973 | 2,238899461 |
| **mmu-miR-7688-5p** | -1,833452 | 2,044986097 |
| **mmu-miR-23b-3p** | 1,638435 | 1,999153682 |
| **mmu-miR-708-5p** | 3,428667 | 1,945652747 |
| **mmu-miR-193a-5p** | -1,780358 | 1,78195532 |
| **mmu-miR-1983** | -1,541618 | 1,774716187 |
| **mmu-miR-221-5p** | 1,929952 | 1,804894565 |
| **mmu-miR-30c-1-3p** | -1,605333 | 1,695938101 |
| **mmu-miR-505-3p** | 5,45504 | 1,701438448 |
| **mmu-miR-669c-5p** | -1,586156 | 1,839462837 |
| **mmu-miR-6998-3p** | 4,461104 | 1,691174154 |
| **mmu-miR-7054-5p** | -2,892144 | 1,750329155 |
| **mmu-miR-9769-3p** | 11,22336 | 1,745859394 |
| **mmu-miR-99a-5p** | 1,635672 | 1,779719382 |
| **mmu-miR-466a-5p** | -6,25811 | 1,588970959 |
| **mmu-miR-6239** | 4,380789 | 1,6045598 |
| **mmu-miR-335-5p** | -2,749834 | 1,558512104 |
| **mmu-miR-1a-3p** | -3,10324 | 1,526204828 |
| **mmu-miR-21a-3p** | 2,957127 | 1,461707432 |
| **mmu-miR-345-5p** | 1,993467 | 1,516059167 |
| **mmu-miR-365-3p** | 1,807271 | 1,503201898 |
| **mmu-miR-501-5p** | 8,532644 | 1,463063319 |
| **mmu-miR-7024-5p** | -5,097571 | 1,476120889 |
| **mmu-miR-204-5p** | -8,006198 | 1,412569502 |
| **mmu-miR-139-3p** | -1,638413 | 1,37398831 |
| **mmu-miR-1947-5p** | -3,44084 | 1,374378562 |
| **mmu-miR-25-5p** | -1,545388 | 1,358440872 |
| **mmu-miR-23b-5p** | -2,424577 | 1,332006406 |
| **mmu-miR-669e-5p** | 4,028704 | 1,341378598 |

The 38 miRNAs are ranked by p-value.

#### mRNA targets and canonical pathway analysis

Since miRNAs carried by EVs can be transferred to recipient cells and mediate the latter’s response, we sought to identify brain cell mRNAs targeted by the four upregulated miRNAs in MPS-EVs. In an expression pairing analysis using the mRNA target filter from Ingenuity Pathway Analysis (IPA), we found 130 predicted mRNAs targets for the four miRNA outliers. To further investigate the pathways in which these 130 mRNA targets were potentially involved, we performed a canonical pathway analysis. The 10 most significant canonical pathways identified without any filter are shown in Figure 4A. The *Neuroinflammation*, *Hepatic fibrosis* and *Role of macrophage* pathways had the highest number of genes, and *Myelination* was ranked fifth. Since mRNA expression vary considerably from one cell type to another, we used *Neurotransmitter and other nervous system signaling* as a filter and found that the top five canonical pathways were associated with *Neuroinflammation*, *Myelination*, *Axonal guidance*, *Huntington’s disease* and *Synaptogenesis* signaling pathways (Fig. 4B). miR-146a-5p and miR-100-5p mainly targeted genes involved in neuroinflammation and myelination, respectively (Fig. 4C), miR-221-3p mostly targeted genes involved in neuroinflammation and axonal guidance signaling pathways, while miR-155-5p targeted genes regulating neuroinflammation and myelination.

**Fig. 4.**
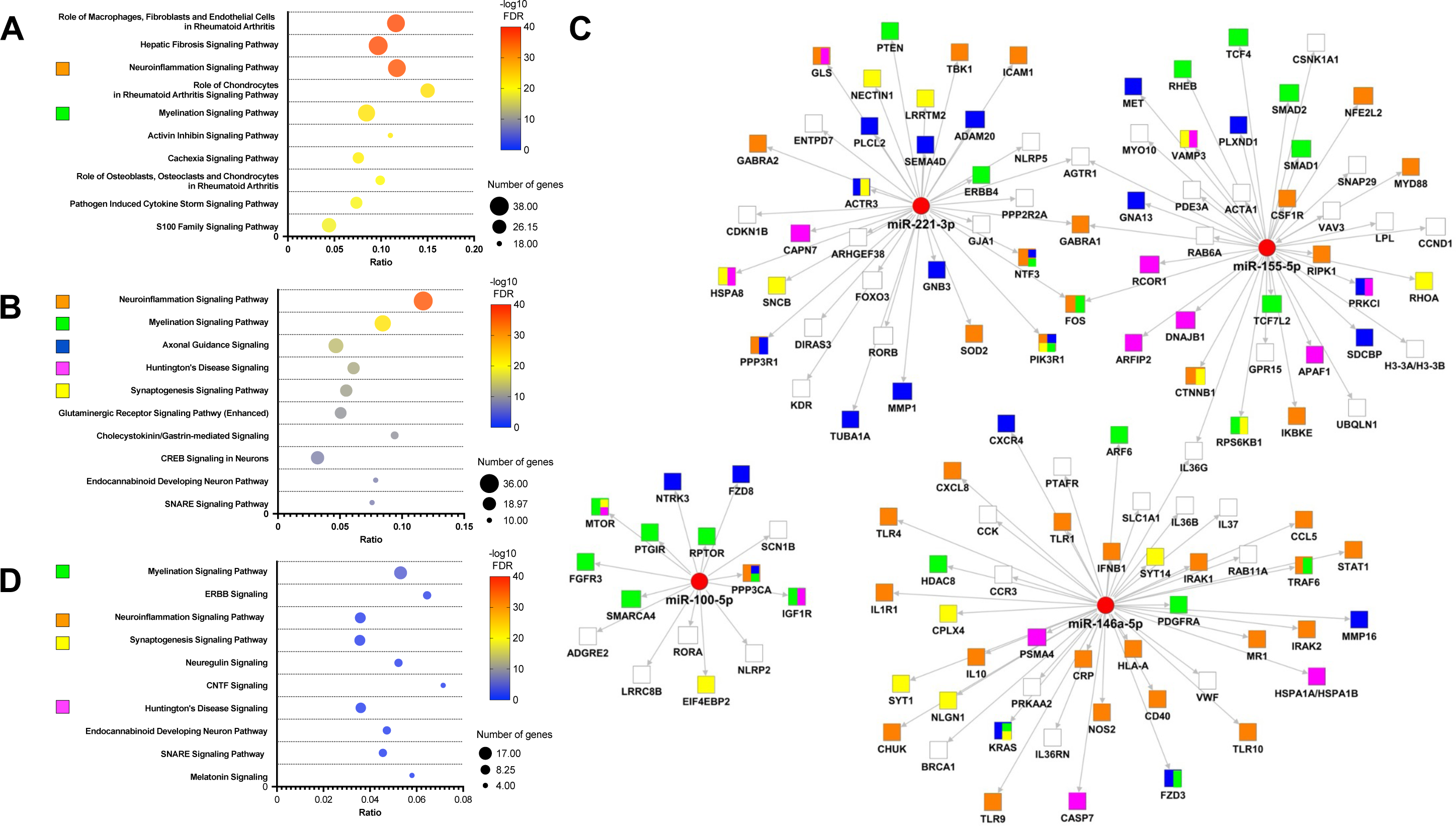
Functional enrichment of mRNAs targeted by the four miRNAs enriched in MPS III-EVs. **(A)** The top 10 significant canonical pathways identified without a filter; the majority were related to inflammatory processes and two pathways specific for neuronal signaling. **(B)** The top 10 significant canonical pathways identified with the *Nervous system* filter, revealing pathways involved in neuroinflammation, neurodevelopment, and neurotransmission. **(C)** A network hub representation of mRNAs targeted by the four EV-miRNAs. Each mRNA (shown as squares) is linked to a miRNA (shown as red circles) with an arrow. The top five IPA canonical pathways with the *Nervous system* filter are represented by colors in each mRNA squares. **(D)** The top 10 significant canonical pathways identified with the *Nervous system* filter for downregulated mRNAs from the transcriptomic analysis of hippocampi of the MPS IIIC *Hgsnat^P304L^* mouse model and that are targeted by the four miRNAs, **(A, B and D)** IPA canonical pathways were ranked by their –log10 FDR. Each pathway is shown as a circle, with the color representing –log10 FDR. The size corresponds to the number of targeted genes, and the X axis represents the gene ratios. The color code for signaling pathways is as follows: Neuroinflammation in orange, Myelination in green, Axonal Guidance in blue, Huntington’s Disease in pink, and Synaptogenesis in yellow.

Using the public mRNA differential expression dataset from the hippocampus of the mouse model of MPS IIIC (21), we identified a total of 648 genes targeted by the four upregulated miRNAs. A canonical pathway analysis showed that the 341 downregulated genes were involved in pathways also identified by prediction analysis: *Myelination*, *Neuroinflammation*, *Synaptogenesis*, *Huntington’s disease*, *Endocannabinoid developing neuron* and *SNARE* signaling pathways (Fig. 4D). Sixty-four of the downregulated mRNAs were also found in the list of 130 genes generated by the prediction analysis.

### Effects of MPS III-EV uptake by primary cortical neurons

#### EV treatment does not reduce neuronal survival

To further explore the neuronal reaction to the EV cargo revealed by our omics experiments, we exposed primary cortical neurons isolated from WT pups to WT-EVs, MPS-EVs or CTL-EVs (8.75x10^8^ EVs/mL) every 24 hours for 72 hours (Fig. 5A). As EVs from the 12k and 100k pellets had similar protein and RNA contents (when considering differences due to GAGs treatment), the two were pooled for the following experiments. We first checked that the number of neuronal nuclei did not change from one condition to another or over time during the treatments (Additional file 1: Fig. S6A). An analysis of TUNEL+ and Hoechst+ colocalization showed that exposure to the EVs did not induce apoptosis of the neurons (Additional file 1: Fig. S6B).

**Fig. 5.**
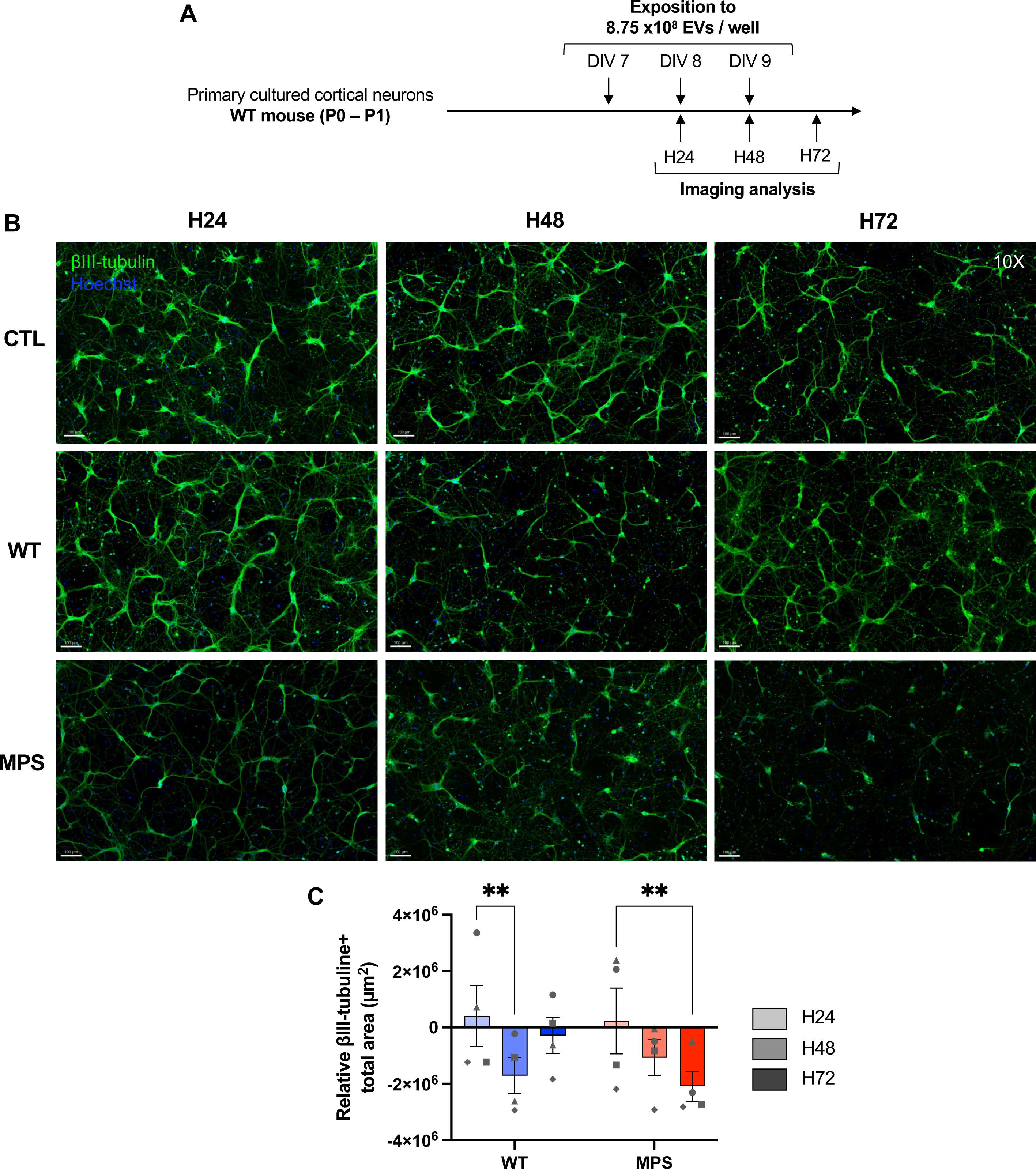
βIII-tubulin immunostaining of primary cortical neurons treated with WT- or MPS-EVs. **(A)** A diagram showing the experiment scheme for chronic exposure of neurons to EVs and the time points in the immunofluorescence imaging analysis. **(B)** Representative wide-field zoomed images of 25-square stitched mosaics (10X objective) showed that βIII-tubulin immunostaining was less intense after neurons had been treated for 72h with MPS-EVs, relative to treatment with WT-EVS or CTL. **(C)** The total βIII-tubulin+ surface area decreased over time in neurons exposed to MPS-EVs. Values for neurons treated with WT-EVs (blue) or MPS-EVs (red) were normalized against those for control neurons (CTL). Two-way ANOVA, n=4 biological replicates (each point represents 1 n), ** = p ≤ 0.01. **(A-B)** Neurons were treated with WT-EVs (WT), MPS-EVs (MPS) or with trehalose (CTL) for 24h (H24, in light grey), 48h (H48, in medium grey) or 72h (H72, dark grey). Neurons were stained with an anti-βIII-tubulin antibody coupled to Alexa Fluor^TM^ 647 (green) and nuclei were stained with Hoechst 33342 (blue). Scale bar = 100 µm.

#### EVs secreted by microglia treated with MPS-GAGs reduce the neurite surface area

Analysis of large mosaic images (25 squares, 10x objective) using Imaris machine-learning software enabled us to quantify the dimensions of the neuronal extensions (Fig. 5B). After normalization against control condition, neuron-specific microtubule immunostaining with an anti-βIII-tubulin antibody highlighted a significant decrease over time in the neurite surface area of neurons exposed to MPS-EVs. The decrease became statistically significant after 72h (two-way ANOVA, p=0.0056, n=4, Fig. 5C). The neurite surface area had decreased significantly in neurons after 48h of exposure to WT-EVs (two-way ANOVA, p=0.0097, n=4) but returned to the baseline value at 72h (Fig. 5C).

#### EVs secreted by microglia treated with MPS-GAGs impair dendritic organization

We investigated the site of neurite damage by immunostaining the dendrites of postmitotic neurons with an antibody against microtubule-associated protein 2 (MAP2) (Fig. 6A). The MAP2+ area in neurons exposed to MPS-EVs decreased over time, and the decrease became statistically significant after 72h (two-way ANOVA, p=0.0281, n=4, Fig. 6B). For neurons treated with WT-EVs, the dendrite surface area decreased between 24h and 48h (two-way ANOVA, p=0.0039, n=4) but returned to the baseline value at 72h (two-way ANOVA, p=0.0369, n=4, Fig. 6B). The MPS-EVs had a clear effect on dendrite length (i.e. the distance from the soma), with a significant decrease in MAP2+ length between 48 h and 72 h (two-way ANOVA, p=0.0369, n=4) and particularly between 24 h and 72 h (two-way ANOVA, p=0.0058, n=4, Fig. 6C).

**Fig. 6.**
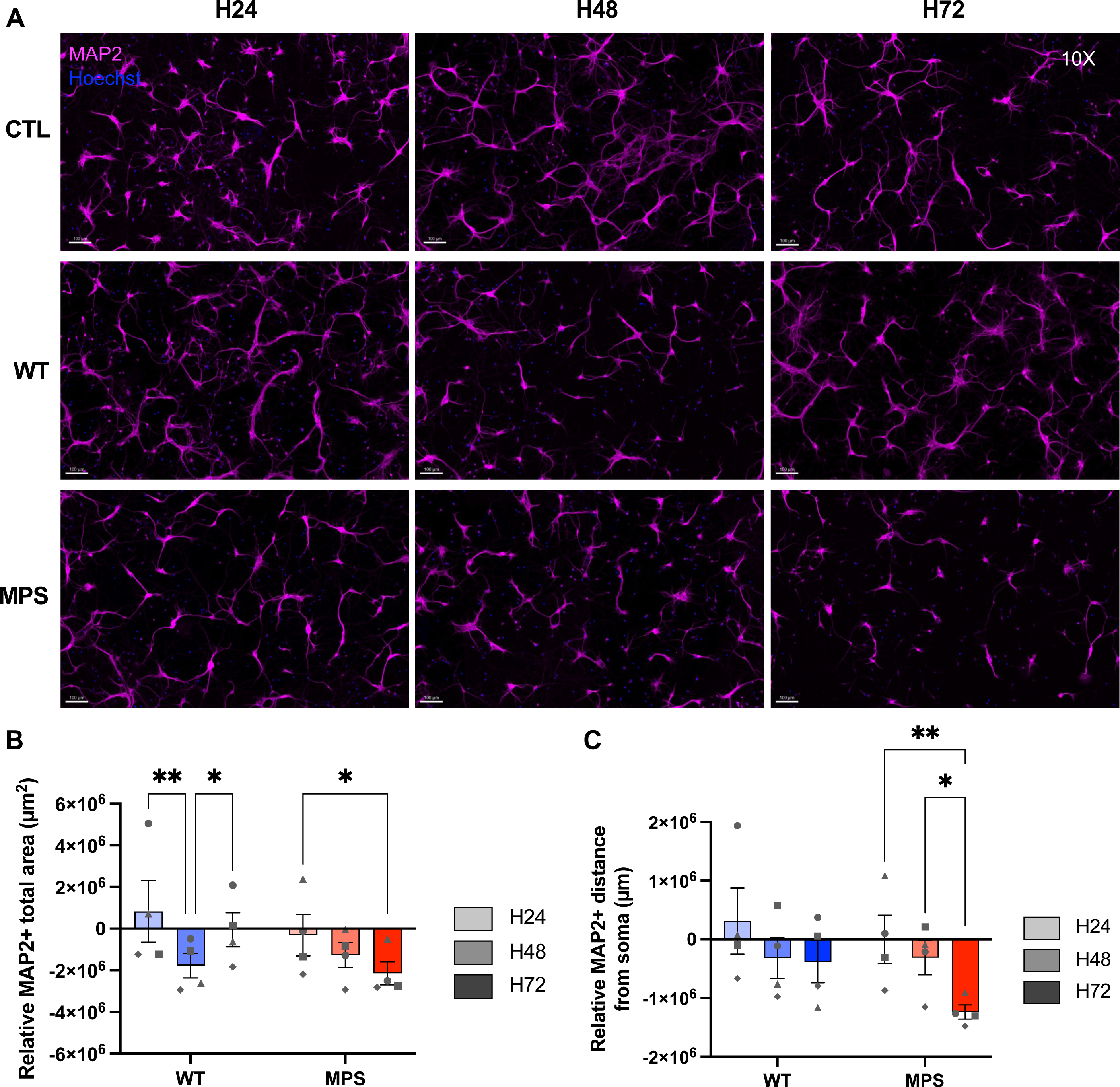
MAP2 immunostaining of primary cortical neurons treated with WT- or MPS-EVs. **(A)** Representative insets of wide-field zoomed images of neurons on 25 squares mosaics (10X objective) showed a relative decrease in the intensity of MAP2 immunostaining in neurons after 72h of treatment with MPS-EVs. **(B)** Quantification of the total MAP2+ surface area revealed a time-dependent decrease in neurons treated with MPS-EVs. **(C)** Quantification of the MAP2+ distance from the soma (dendrite length) revealed a progressive decrease in neurons treated with MPS-EVs. **(A-C)** Neurons were treated with WT-EVs (WT, blue), MPS-EVs (MPS, red) or trehalose (CTL) for 24h (H24, in light grey), 48h (H48, in medium grey) or 72h (H72, in dark grey). Neurons were stained with an anti-MAP2 antibody coupled to Alexa Fluor^TM^ 568 (in pink) and nuclei were stained with Hoechst 33342 (in blue). Scale bar = 100 µm **(B and C)** Quantifications of wide-field neurons variables, normalized against nontreated control neurons (CTL). Two-way ANOVA, n=4 biological replicates (each point represents 1 n), * = p ≤ 0.05; ** = p ≤ 0.01; *** = p ≤ 0.001; **** = p ≤ 0.0001.

We also exposed neurons to EVs secreted by microglia activated for 24 h with 100 ng/mL of LPS (LPS-EVs) in order to investigate whether the inflammatory component alone is responsible for neuronal network disorganization. After validating by RT-qPCR the enrichment of the inflammatory miR-146a-5p and miR-155-5p in LPS-EVs, we showed that after 72 h of treatment, there was no significant decrease in βIII-tubulin+ surface area or in MAP2+ surface area and length (data not shown, Additional file 3).

Under the microscope (40X objective, Fig. 7A), neurons exposed to MPS-EVs showed an increase in the number of short MAP2+ protrusions from the soma; the increase was statistically significant at 48 h and at 72 h (two-way ANOVA, n=4, p=0.0263 and p=0.0034, respectively) (Fig. 7B). The increase in the number of MAP2+ branches after exposure to MPS-EVs was also significant when compared with neurons treated with WT-EVs at 48 h and at 72 h (two-way ANOVA, n=4, p=0.0039 and p=0.0026, respectively) (Fig. 7B). At the 72 h timepoint, we observed an increase in the number of short primary MAP2+ branches (two-way ANOVA, p=0.0016, n=4, Fig. 7C) but not in second-generation branches. However, we observed a decrease in the number of tertiary (terminal) branches (two-way ANOVA, p=0.0022, n=4, Fig. 7C).

**Fig. 7.**
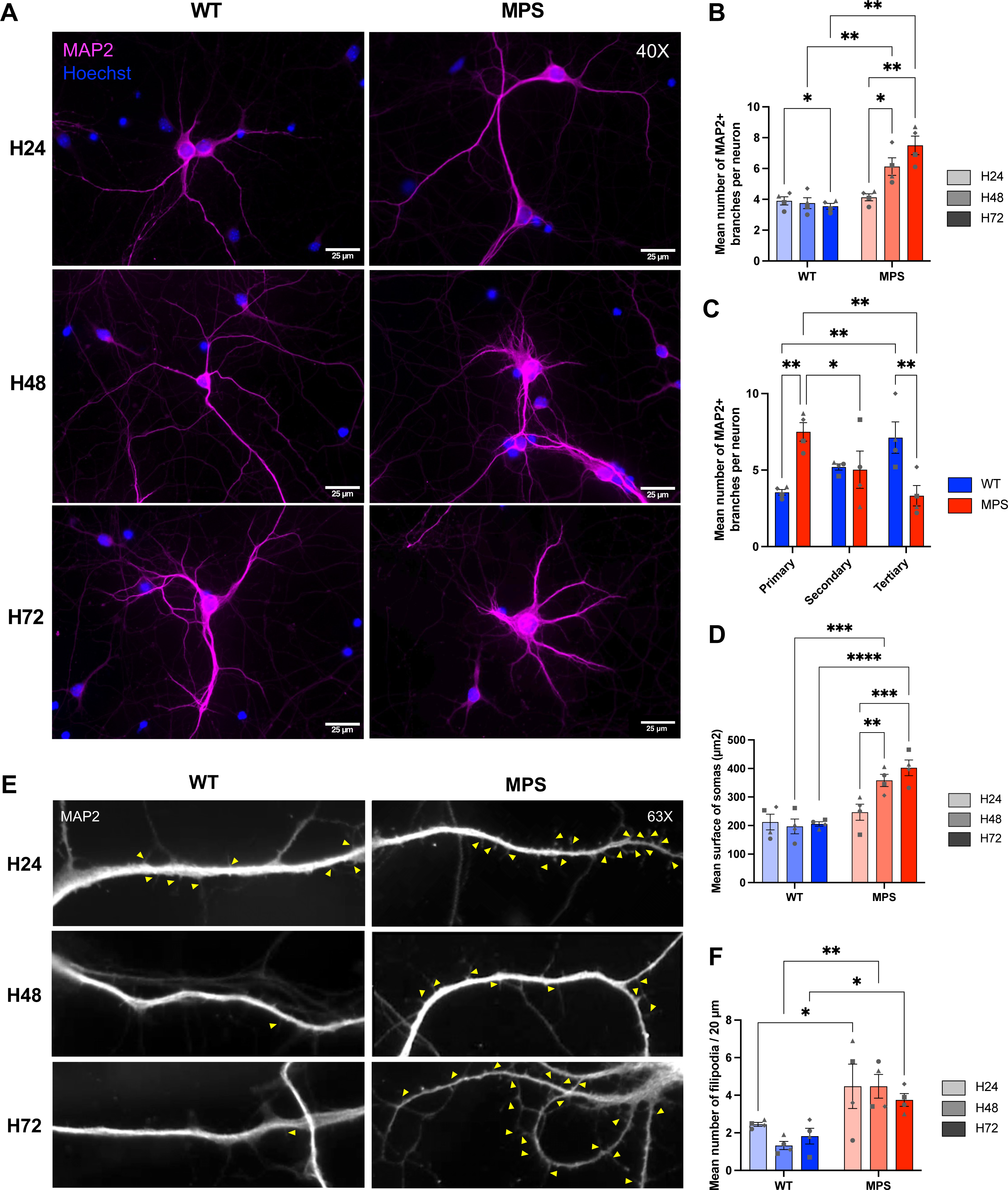
Characterization of MAP2+ dendrites on primary cortical neurons treated with WT- or MPS-EVs. **(A)** Representative images of 1 or 2 neurons (40X objective). **(B)** The number of short MAP2+ dendritic branches increased progressively in the MPS condition. **(C)** Quantification of the number of primary and tertiary branches per neurons revealed an overall decrease in the complexity of the dendritic arborization in neurons treated with MPS-EVs after 72h. **(D)** Quantification of the soma’s surface area revealed the progressive swelling of neurons treated with MPS-EVs. **(B-D)** Quantification of somatodendritic variables from 10 neurons per n. **(E)** Representative images of dendrites (63X objective) with filipodia spines (>2 μm, yellow arrowheads), showing an increase the number of filipodia per dendrite in neurons treated with MPS-EVs. **(F)** The number of immature dendritic spines increased in neurons treated with MPS-EVs. The filipodia were analyzed 30 μm away from the soma on 20-μm-long MAP2+ segments. Data from 10 dendrites from 5-10 neurons were expressed as the mean value per n. **(A-F)** Neurons were treated with WT-EVs (WT, in blue) or MPS-EVs (MPS, in red) for 24h (H24, in light grey), 48h (H48, in medium grey) or 72h (H72, in dark grey). Neurons were stained with an anti-MAP2 antibody coupled to Alexa Fluor^TM^ 568 (in pink in A and in white in E) and nuclei were stained with Hoechst 33342 (in blue). Scale bar = 25 µm (A), 20 µm (E). Two-way ANOVA, n=4 biological replicates (each point represents 1 n), * = p ≤ 0.05; ** = p ≤ 0.01; *** = p ≤ 0.001; **** = p ≤ 0.0001.

We also observed soma swelling after MPS-EV treatment, as evidenced by an increase in the soma area surface at 48 h and at 72 h (two-way ANOVA, n=4, p=0.0054, and p=0.0005, respectively). This difference in surface area was also statistically significative in neurons treated with MPS-EVs vs. WT-EVs at 48 h and at 72 h (two-way ANOVA, n=4, p=0.0002 and p<0.0001, respectively; Fig. 7D).

Lastly, we counted the number of immature long dendritic spines (i.e. filipodia) per 20 µm of dendrite (63X objective). Some filipodia were more than 2 µm long (yellow arrow heads in Fig. 7E). The number of MAP2+ filipodia on neurons exposed to MPS-EVs increased at 24 h, 48 h and 72 h (two-way ANOVA, n=4, p=0.0273, p=0.0015 and p=0.0348, respectively; Fig. 7F).

## Discussion

Severe neuroinflammation is a hallmark of MPS III in both patients and mouse models. Chronic microglial activation has been shown to have a prominent role in the neuroinflammatory process that contributes to progressive neurodegeneration. The impact of EVs released by microglia on the physiopathology of MPS III has not been investigated.

It is well known that the BV-2 microglia murine cell line can be stimulated by lipopolysaccharide (LPS) and secretes pro-inflammatory cytokines that modulate oxidative stress or apoptosis (47,48). Hence, the BV-2 cell line is the gold standard in neuroinflammation studies (49) and studies of neurodegenerative disorders (50). Here, we used GAGs purified from the urine of patients with MPS III. We used this GAG fraction (highly enriched in undigested HS) to activate microglia and study their EVs. We have shown previously that HSOs have damaging effects on neurons, astrocytes, and microglia (10,17,51–53).

### MPS III microglia-derived EVs carry a cargo of proteins and miRNAs involved in the inflammatory response and neuronal development

We first characterized the RNAs and proteins in EVs released by microglial cells treated with GAGs isolated from the urine of children with MPS III. Few researchers have used proteomics or multi-omics analyses to comprehensively characterized the molecular signature of microglial EVs produced in neurodegenerative states. In exosomes produced in TREM2 Alzheimer’s disease (AD) risk variant iPS-microglia, Mallach et al. found nine enriched proteins involved in the downregulation of transcription and metabolic processes (54). In CD11b^+^ small EVs from the brains of people with AD, 27 proteins were differently expressed. These proteins included disease-associated microglia markers (TMEM119, P2RY12, FTH1, ApoE) and neuronal and synaptic proteins (55). Mallach et al. also identified four miRNAs (miR-28-5p, -381-3p, -651-5p, and -188-5p) that were more abundant in microglial EVs from people with AD. These miRNAs regulated the Toll-like receptor (TLR), and senescence pathways.

The results of our RNA sequencing experiments showed that three miRNAs (-146a-5p, -155-5p and –221-3p, all known to be involved in the immune response) were abundant in MPS III microglial-EVs. Interestingly, both miR-146a-5p and miR-155-5p were found to be upregulated in EVs secreted by LPS-polarized N9 microglia (56), miR-146-5p in Aβ-treated microglial EVs (29), and miR-221-3p in EVs from IL-4 polarized microglia (57); these observations indicated that in an inflammatory context, microglia release EVs with a high abundance of these specific miRNAs.

Similarly, comparative proteomic analyses revealed that all the proteins enriched in MPS III-EVs were involved in the inflammatory response or immune cell recruitment. In particular, some of these proteins were part of the *Toll-like receptor signaling, and lipopeptide binding* PPI network. This inflammatory protein and miRNA cargo is consistent with the reported ability of

HS and HSOs to promote an innate immune response. The induction of the pro-inflammatory cytokine release by HS fragments (via interactions with TLR4) has been observed under physiological conditions *in vitro* (58,59). Furthermore, we have shown previously that HSOs can activate microglia through the TLR4/MyD88 pathway (10,17). In both Naglu/Tlr4 and Naglu/Myd88 double-knockout mice, the onset of brain inflammation was delayed for several months when compared with MPS IIIB mice. In MPS VII, inactivation of TLR4 pathway in GusB/Tlr4 double-knock-out mice corrects many biochemical and clinical features of the disease (60). However, we demonstrated that even if MPS-EVs are enriched in inflammatory content, an exclusive pro-inflammatory signature (e.g. LPS-EVs) is not sufficient to initiate neurites disorganization.

Interestingly, most of the proteins that were less abundant in MPS III microglial EVs turned out to be involved in neuronal development mechanisms, axonogenesis, gliogenesis, or dendritic spine morphogenesis. Furthermore, a PPI network analysis identified the *Axon guidance and basigin-like* and *Toll-like signaling, and lipopeptide binding* protein networks. It is widely acknowledged that miRNA expression is inversely related to target gene expression. In the light of the results from our expression pairing analysis, we looked at whether the predicted mRNA targets were truly downregulated *in vivo* by using the published differential mRNA expression dataset for the hippocampus of the mouse model of MPS III C (21). The resulting canonical pathway analysis of genes targeted by the four miRNAs revealed that *Myelination*, *Huntington’s disease* and *Synaptogenesis* were the top pathways in terms of both predicted mRNA targets and the transcriptomic analysis of MPS IIIC hippocampi. It is notable that miR-100-5p (which regulates genes closely involved in neuronal development) has not previously been found to be upregulated in EVs secreted by microglia. Interestingly, Li et al. reported that following spinal cord injury, miR-100 attenuates the inflammatory activation of microglia (by inhibiting the TLR4/NF-κB pathway and neuronal tissue apoptosis) and improves motor function (61).

Our comprehensive characterization of the cargo of microglia-derived EVs (by combining proteomics and transcriptomics) reflected the microglia’s multiple functions as a CNS immune cell and as a support for neurons. Moreover, our results suggest that these two roles are altered in MPS III-microglia: the latter’s small and large EVs are enriched in proteins with a role in inflammatory processes and lack proteins involved in neuronal development. Concordantly, the abundant miRNAs in these EVs can regulate genes involved in the same biological pathways.

We have shown previously that inactivation of the TLR4-MyD88 pathway in MPS IIIB mice suppressed early-onset neuroinflammation but did not prevent neurologic disease (10). We therefore hypothesize that microglial EVs that lack neurotrophic molecules have a role in the progression of neurodegeneration.

According to the literature data, this molecular profile appears to be very specific for EVs secreted by MPS III microglia. Hence, the cargo might be harmful for the recipient cells. This was further demonstrated by treating primary cortical neurons with EVs secreted by microglia exposed to HSOs isolated from the urine of patients with MPS III.

### MPS III microglia-derived EVs impair dendrite arborization in primary cortical neurons

When immunostaining the neuronal cytoskeleton and the somatodendritic compartment with anti-βIII-tubulin and anti-MAP2 antibodies respectively, we observed significant, similar decreases over time in the numbers of neurites and dendrites in neurons treated with MPS III-microglia-EVs (MPS III-Mg-EVs). We next characterized the dendrites’ arborization. Interestingly, we found that neurons treated with MPS III-Mg-EVs presented fewer tertiary branches and more primary dendrites. Furthermore, the primary dendrites were shorter (i.e. closer to soma). These experiments demonstrated that exposure to MPS III-Mg-EVs was associated with a less complex arborization – mainly due to fewer tertiary branches. Interestingly, similar observations have been made in models of MPS III. Hocquemiller et al. reported defects in dendritic arborization in live primary cortical neurons of MPS IIIB mice (52). More recently, a decrease in neurites length was also reported in primary hippocampal neurons of MPS IIIC mice. Pará et al. also showed that in pyramidal neurons, the number of dendritic spines was low while the number of immature dendritic spines was high. We also observed that primary cortical neurons treated with MPS III-Mg-EVs had a greater number of immature dendritic spines, suggesting that a low number of synapses could impair neuronal transmission. Indeed, MPS IIIC neurons presented early abnormalities in synaptic structure and neurotransmission as a result of impaired synaptic vesicular transport (62).

A time-dependent increase in the soma surface area (described in the literature as “swelling” or “ballooning”) was observed in neurons treated with MPS III-Mg-EVs. This is a histological feature of neurodegeneration and has been reported in tauopathies. For example, the accumulation of abnormal cytoskeletal components (e.g. the formation of neurofibrillary tangles in AD and the formation of Lewy bodies) can cause swelling of the cell body. In metabolic diseases, accumulation of storage material reportedly induce neuron swelling, and swollen neurons have been reported in the brains of patients with MPS IIIB (63) and in the canine model of MPS IIIB (64).

Under pathological conditions, the uptake of microglia-derived EVs reportedly impairs neuronal functions. Recently, Prada et al. showed *in vitro* that EVs from pro-inflammatory microglia can affect neuronal transmission, with a lower density of mature dendritic spines (29). Small EVs from TREM2 AD risk variant iPS-microglia were less able to promote the outgrowth of neuronal processes (30). The striatal injection of exosomes isolated from α-synuclein-treated microglia induced neurodegeneration in the nigrostriatal pathway of injected mice (26). Furthermore, EVs derived from the brains of patients with AD are known to transport tau proteins and mediate the dysfunction of GABAergic interneurons (27,65). In these studies, the effect of EVs was mainly attributed to the transmission of the abnormal protein to the neuron. We cannot attribute the effects observed in our study to the transport of HSO by EVs; in the vitro model used here, the HSOs purified from the urine of patients with MPS III were added extracellularly and were unlikely to accumulate in the way seen in the mouse models of MPS III. It would be therefore very interesting to determine *in vivo* whether MPS III-Mg-EVs transport HS fragments. This could be measured using a very sensitive LC-MS/MS system.

## Conclusions

Our results showed that the characteristic cargo of MPS III-Mg-EVs can induce a MPS III-like phenotype in the recipient naive neurons - probably through the impairment of neurotransmission. Indeed, our results strongly suggest that in patients with MPS III, microglia-derived EVs accentuate inflammation and neuronal dysfunction. The present study is the first to have characterized the content of MPS III-Mg-EVs. We observed a disease-associated signature and provided a framework for further studies of patient-derived, cell-type-specific EVs and their potential value as biomarkers of the response to treatment of MPS III.

## List of abbreviations

AD: Alzheimer’s disease
ADAM10: A Disintegrin and metalloproteinase domain-containing protein 10
CNS: Central nervous system
CPM: count per million
EV: Extracellular Vesicles
FDR: False discovery rate
GAGs: Glycosaminoglycans
HSOs: Heparan Sulfate Oligosaccharides
HSPGs: Heparan Sulfate Proteoglycans
IPA: Ingenuity Pathway analysis
Lamp1: Lysosomal Associated Membrane Protein 1
LC-MS/MS: Liquid Chromatography coupled to tandem Mass Spectrometry
LSD: Lysosomal Storage Disease
MAP2: microtubule-associated protein 2
miRNA: microRNA
MPS III: Mucopolysaccharidosis type III
MyD88: Myeloid differentiation primary response 88
NTA: Nanoparticle tracking analysis
PPI: protein-protein interaction
SEM: standard error of the mean
STRING: Search Tool for the Retrieval of Interacting Genes/Proteins
TEM: Transmission Electron microscopy
TfR: Transferrin receptor
TLR: Toll-like receptor
TUNEL: Terminal deoxynucleotidyl transferase dUTP nick end labeling
WT: Wild type

## Declarations

### Ethics approval and consent to participate

In line with the French Public Health Code (*Code de la Santé Publique*, article L1211-2), the parents and/or their legal guardians of under-18 MPSIII patients and of under-18 healthy control volunteers stated that they did not object to the use of urine samples for biomedical research. Mouse experiments were performed by authorized investigators in accordance with French regulations and the European Union Council Directive 86/609/EEC and were approved by the government-accredited animal care and use committee (*Comité d’Ethique de l’US 006/CREFRE* (Toulouse, France); reference number: PI-U1291-JA-37).

### Consent for publication

Not applicable

### Availability of data and materials

The mass spectrometry proteomics data have been deposited to the ProteomeXchange Consortium via the PRIDE partner repository (http://www.ebi.ac.uk/pride) with the dataset identifier PXD050768. The RNA sequencing data generated in this study are available at Sequence Read Archive website (http://www.ncbi.nlm.nih.gov/bioproject), accession no. PRJNA1090840.

## Funding

This work was funded by the Institut National de la Santé et de la Recherche Médicale (INSERM) and the Vaincre les Maladies Lysosomales (VML) charity.

### Authors’ contributions

CD performed, analyzed, and interpreted the majority of the experiments, conducted the bioinformatic analysis, drafted the manuscript and created the figures. NB carried out the animal experiments, validated the in vitro model, helped with the analysis and interpretation of the data, and created the figures. GM did the isolation of EVs for RNA sequencing and performed and analyzed miRNA experiments. KA performed and analyzed NTA experiments. CS isolated and quantified GAGs from collected urines. CG performed mass spectrometry data analysis and bioinformatics. AC performed RNA sequencing experiments. JA raised the scientific question, conceived the study, and revised the manuscript. ST designed and supervised the study, analyzed, and interpreted the data and wrote the manuscript. All authors read and approved the final manuscript.

## Acknowledgements

We particularly thank our collaborators Alexey V. Pshezhetsky and Mahsa Taherzadeh from CHU Sainte-Justine, Montreal, for sharing their RNAseq dataset. We thank Elsa Suberbielle and Benjamin Schmitt for technical assistance in the isolation of murine primary neurons; Evert Hannapple and Emily Clement for their help in setting up the NTA experiments.

We thank Kévin Roger and Cerina Chhuon at the Necker Proteomics Plateform, Paris, for performing proteomic experiments, bioinformatic and statistical analysis; Margot Zahm and Yann Aubert at Genomic and Transcriptomic platform of Infinity for RNAseq bioinformatic and statistical analysis We also thank for their technical assistance Sophie Allart, Simon Lachambre, and Lhorane Lobjois at the Cellular imaging core facility; Stéphanie Balor and Vanessa Soldan at the Multiscale Electron Microscopy core facility connected to ‘Toulouse Réseau Imagerie’ network; and all the staff from Service Zootechnie Expérimentation Purpan animal core facility.

---

**Fig. S1.**
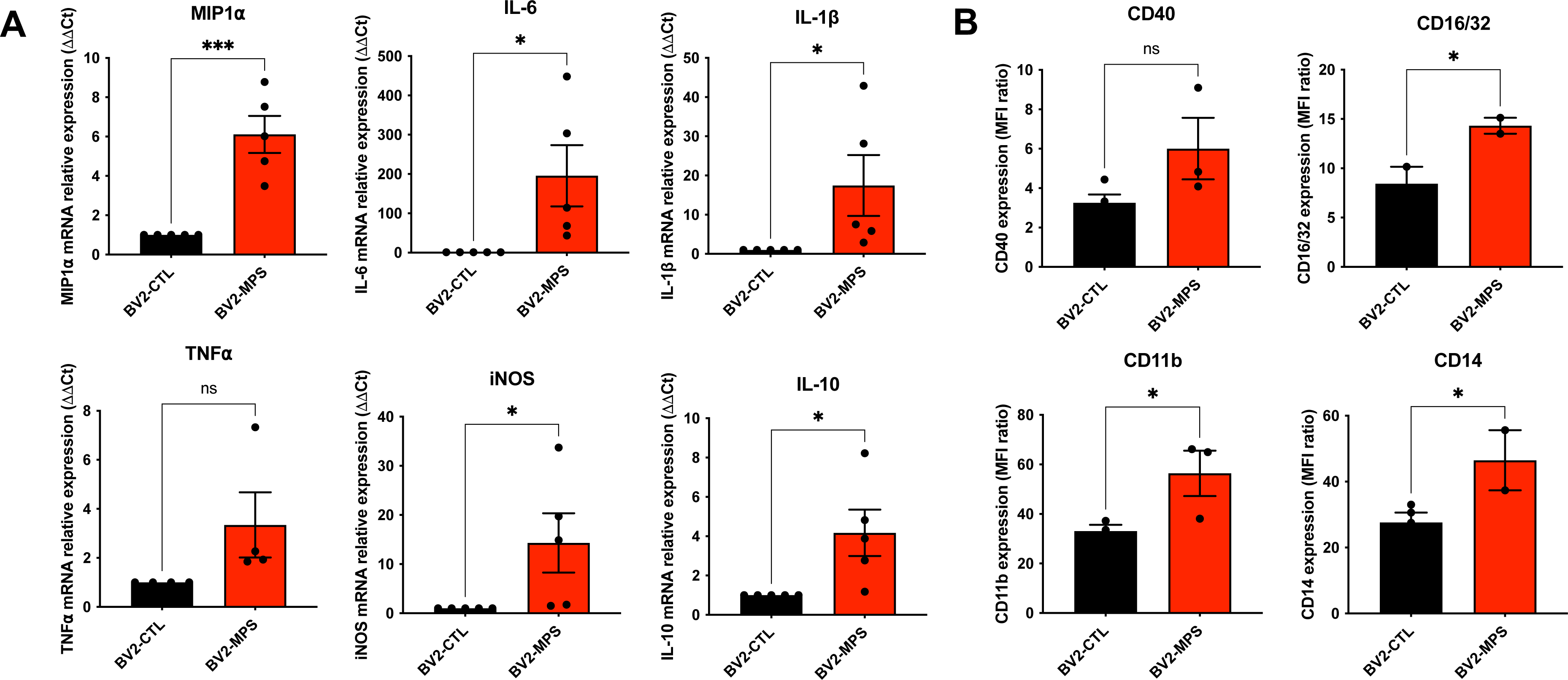
Pro-inflammatory activation of BV-2 cells by GAGs extracted from urine of patients with MPS III. **(A)** An RT-qPCR analysis confirmed higher mRNA expression levels for MIP1⍺ (p=0.0003), IL-6 (p=0.0186), IL-1β (p=0.0335), TNF⍺ (p=0.0643), iNOS (p=0.0291), and IL-10 (p=0.0137) in BV-2-MPS cells. **(B)** Flow cytometry analysis revealed higher protein expression levels for the myeloid surface markers CD16/32 (p=0.0322), CD11b (p=0.0314) and CD14 (p=0.0199) in BV-2-MPS cells. **(A-B)** BV2 cells treated with MPS III GAG (BV2-MPS, in red) were compared with BV-2 cells treated with vehicle only (BV2-CTL, black) in bar plots. Unpaired t-test, n=2-5 biological replicates (each point represents 1 n), one-tailed p-value, ns = p > 0.05; * = p ≤ 0.05; ** = p ≤ 0.01; *** = p ≤ 0.001.

**Fig S2.**
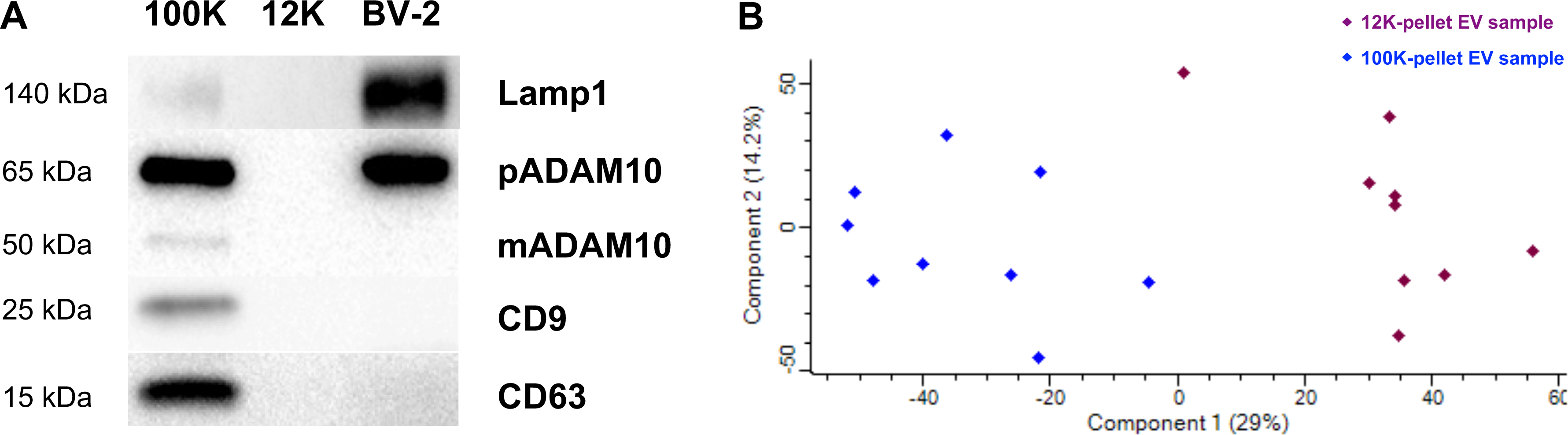
Quality check on proteins from large and small EVs. **(A)** Western blots performed with CTL-100k EVs (100k), CTL-12k EVs (12k) and BV-2 cell lysates (BV-2) revealed the enrichment of specific EV protein markers CD9, CD63, pADAM10 and LAMP1 in CTL-100k EVs. **(B)** The results of a principal component analysis of the EV sample quality check, showing that the 12k-EV population (in brown) and the 100k-EV population (in blue) clustered together.

**Fig. S3.**
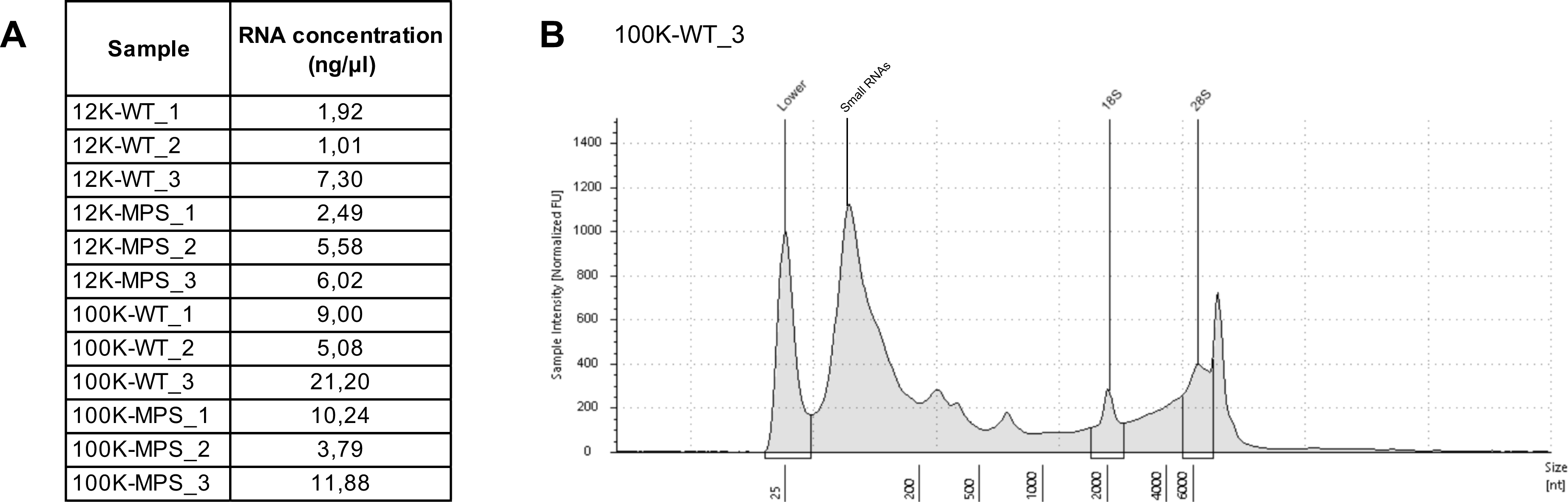
Quality check of RNA samples from large and small EVs. **(A)** Before sequencing, total RNA was quantified using a Qubit Fluorometer (Illumina QIAseq). **(B)** A representative total RNA electrophoresis profile evaluated on an Agilent 4150 TapeStation, showing the presence of small RNAs. Peaks from left to right: lower marker, small RNAs, ribosomal 18S and 28S RNAs.

**Fig. S4.**
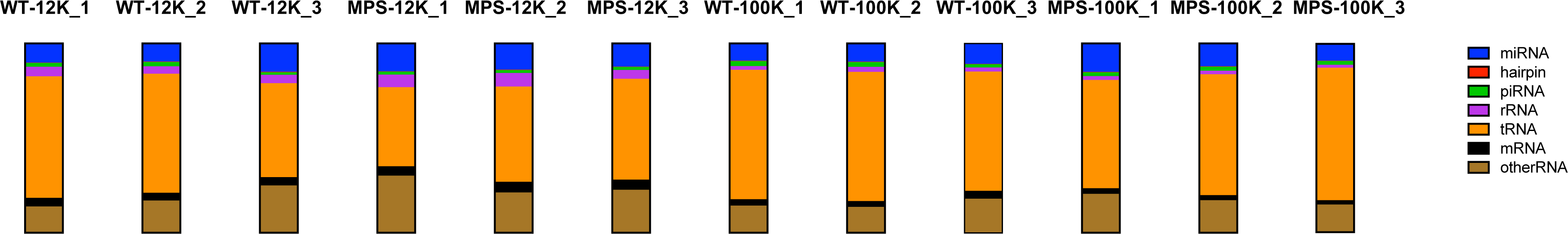
Proportion of different types of RNA in EV samples. Transfer RNAs and miRNAs were the most frequent RNAs. The figure shows miRNAs (in blue), hairpins (in red), piRNAs (in green), rRNAs (in purple), tRNAs (in orange), mRNAs (in black), and other RNAs (in brown).

**Fig. S5.**
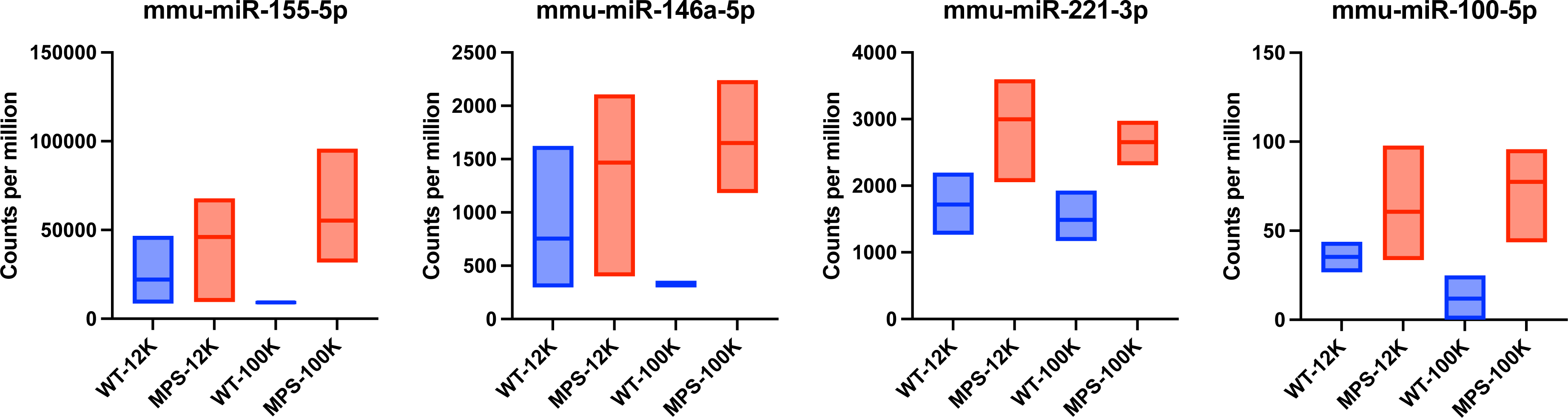
Counts per million for the four miRNAs enriched in MPS-EVs (represented as violin plots). EVs were collected from conditioned media from BV-2 cells treated with WT-GAGs (WT, blue), MPS III-GAGs (MPS, red), and the RNA was extracted from the 12k pellet (12k) or the 100k pellet (100k). n=3 biological replicates (each bar of the violin plot represents 1 n).

**Fig. S6.**
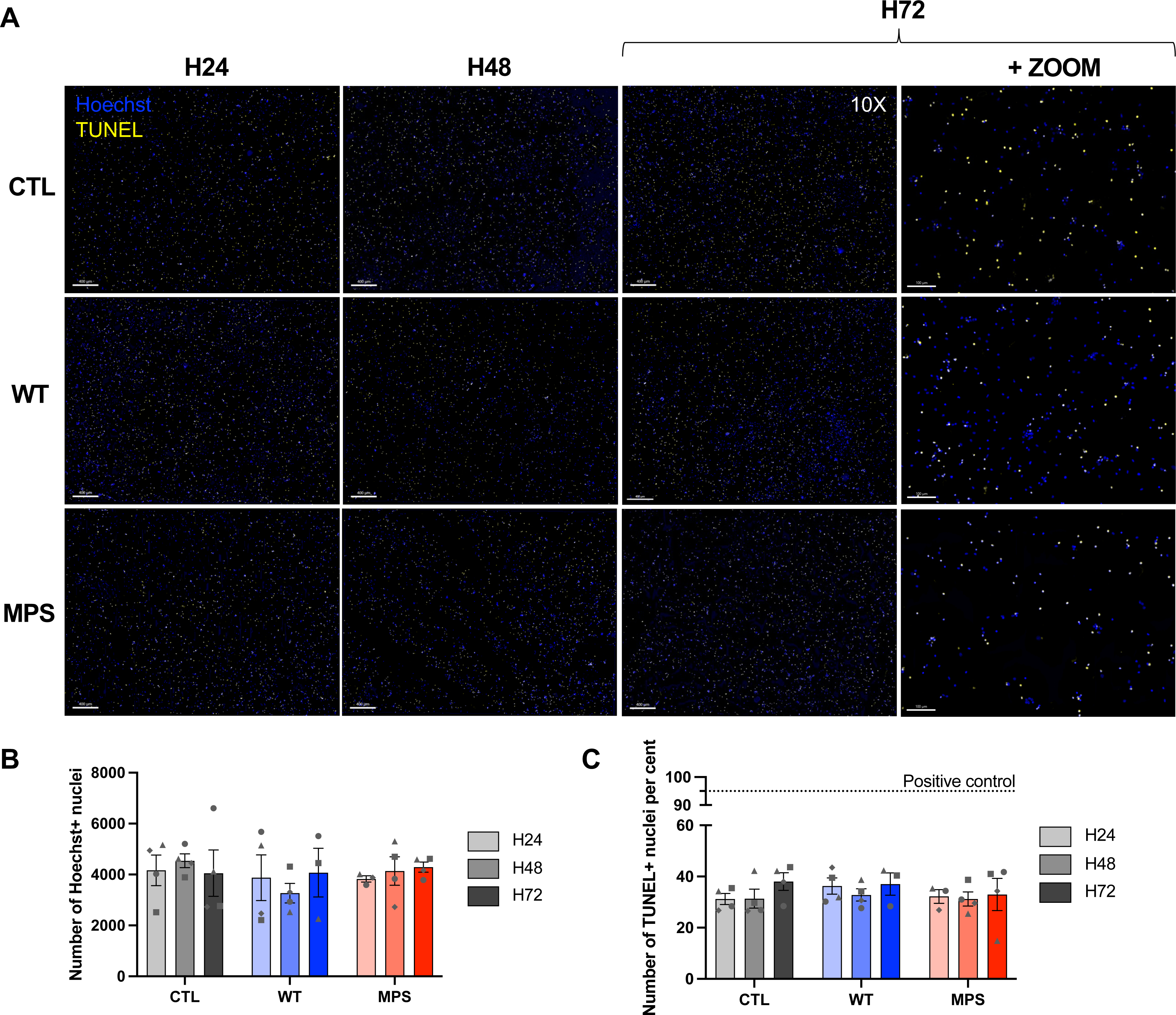
DAPI staining and TUNEL assay results for neurons exposed to EVs. **(A)** Representative wide-field images of 25-square stitched mosaics (10X objective) of neurons treated with WT-EVs (WT), MPS-EVs (MPS) or with trehalose (CTL) for 24 hours (H24), 48 hours (H48) or 72 hours (H72). The nuclei were stained with Hoechst 33342 reagent (in cyan) or Alexa Fluor^TM^ 488 dye (in yellow) when combined with a TUNEL assay. Scale bars = 400 µm for the main images and 100 µm for insets. **(B)** Exposure to EVs did not lead to a decrease in the number of Hoechst+ nuclei. (**C)** Quantification of colocalized TUNEL+ and Hoechst+ nuclei showed that EVs do not have an apoptotic effect on neurons. **(B, C)** Neurons treated with trehalose (CTL, grey), WT-EVs (WT, blue) or MPS-EVs (MPS, red) after 24 hours (H24, in light grey), 48 hours (H48, in medium grey) or 72 hours (H72, in dark grey) of treatment. Two-way ANOVA, n=4 (each point represents 1 n).

**Table S1.**
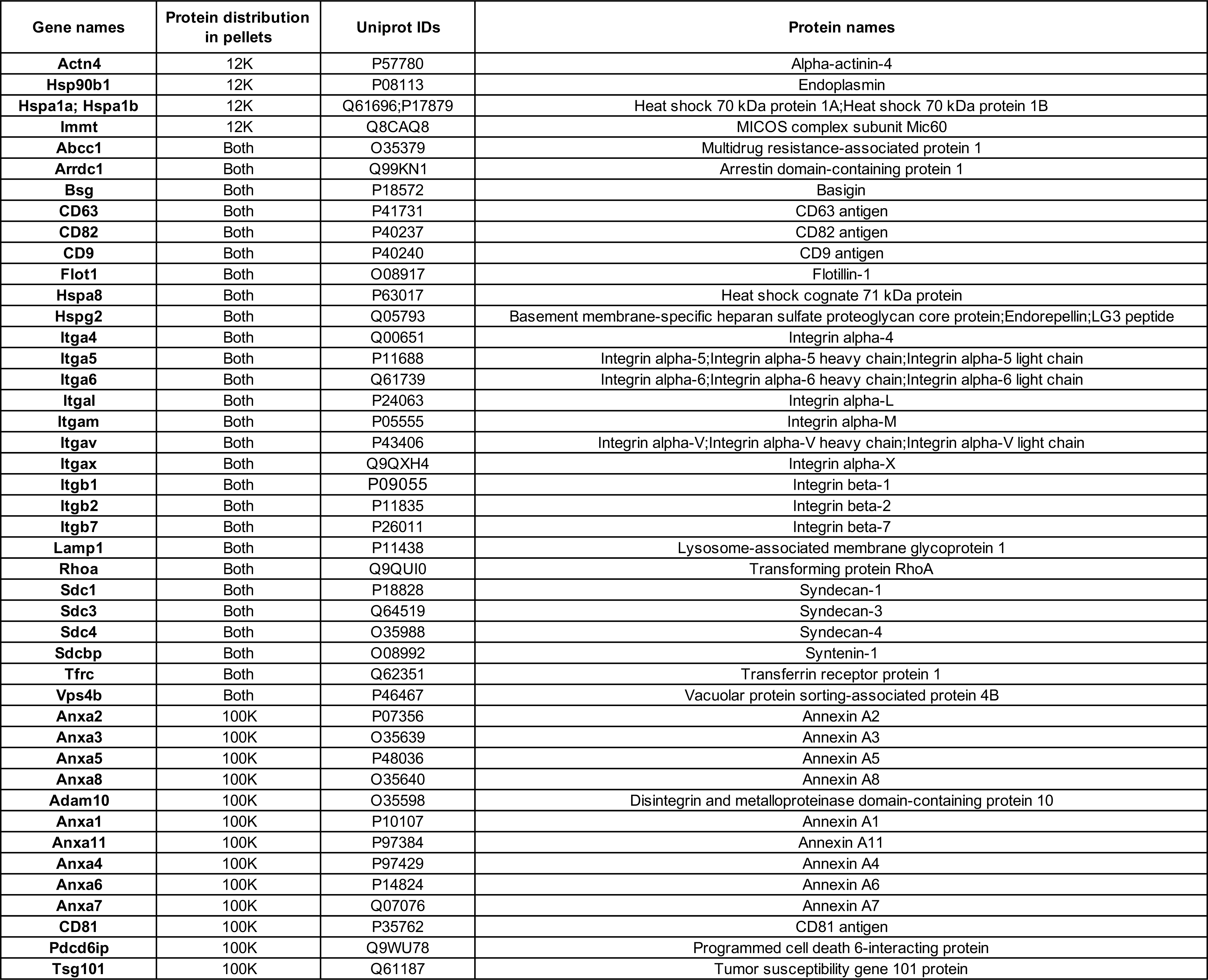
List of specific protein markers of large and small EVs, as identified by mass spectrometry.

**Table S2.** Number of reads and percentage of different types of RNA from each EV sample. The data were produced by the Legacy Analysis Pipeline on the QIAGEN GeneGlobe portal.

| Reads set | 12K-WT_1 |  | 12K-WT_2 |  | 12K-WT_3 |  | 12K-MPS_1 |  | 12K-MPS_2 |  | 12K-MPS_3 |  |
| --- | --- | --- | --- | --- | --- | --- | --- | --- | --- | --- | --- | --- |
|  | Reads number | % | Reads number | % | Reads number | % | Reads number | % | Reads number | % | Reads number | % |
| total_reads | 14 171 376 | 100 | 14 028 393 | 100 | 12 142 987 | 100 | 14 542 659 | 100 | 11 742 528 | 100 | 14 622 539 | 100 |
| no_adapter_reads | 661 606 | 5 | 921 381 | 7 | 1 195 919 | 10 | 750 957 | 5 | 840 160 | 7 | 747 963 | 5 |
| too_short_reads | 2 529 611 | 18 | 3 055 470 | 22 | 1 842 653 | 15 | 2 488 025 | 17 | 1 924 101 | 16 | 2 288 878 | 16 |
| UMI_defective_reads | 914 853 | 6 | 1 277 084 | 9 | 1 865 634 | 15 | 1 625 728 | 11 | 1 397 566 | 12 | 1 731 679 | 12 |
| Total reads - defaults | 10 065 306 | 71 | 8 774 458 | 63 | 7 238 781 | 60 | 9 677 949 | 67 | 7 580 701 | 65 | 9 854 019 | 67 |
| miRNA_Reads | 733 705 | 5 | 523 473 | 4 | 798 777 | 7 | 1 121 081 | 8 | 776 996 | 7 | 869 015 | 6 |
| hairpin_Reads | 11 993 | 0 | 14 180 | 0 | 14 881 | 0 | 14 318 | 0 | 12 703 | 0 | 13 329 | 0 |
| piRNA_Reads | 145 270 | 1 | 121 959 | 1 | 71 946 | 1 | 118 187 | 1 | 86 358 | 1 | 114 866 | 1 |
| rRNA_Reads | 366 310 | 3 | 215 689 | 2 | 234 357 | 2 | 505 560 | 3 | 404 048 | 3 | 326 975 | 2 |
| tRNA_Reads | 4 632 323 | 33 | 3 425 267 | 24 | 2 643 947 | 22 | 3 168 072 | 22 | 2 820 223 | 24 | 3 773 447 | 26 |
| mRNA_Reads | 328 920 | 2 | 225 739 | 2 | 233 529 | 2 | 378 021 | 3 | 316 174 | 3 | 381 509 | 3 |
| otherRNA_Reads | 1 002 687 | 7 | 925 160 | 7 | 1 328 657 | 11 | 2 286 748 | 16 | 1 192 253 | 10 | 1 605 922 | 11 |
| notCharacterized_Mappable | 848 410 | 6 | 948 710 | 7 | 591 278 | 5 | 669 948 | 5 | 617 793 | 5 | 806 159 | 6 |
| notCharacterized_notMappable | 1 995 688 | 14 | 2 374 281 | 17 | 1 321 409 | 11 | 1 416 014 | 10 | 1 354 153 | 12 | 1 962 797 | 13 |

| Reads set | 100K-WT_1 |  | 100K-WT_2 |  | 100K-WT_3 |  | 100K-MPS_1 |  | 100K-MPS_2 |  | 100K-MPS_3 |  |
| --- | --- | --- | --- | --- | --- | --- | --- | --- | --- | --- | --- | --- |
|  | Reads number | % | Reads number | % | Reads number | % | Reads number | % | Reads number | % | Reads number | % |
| total_reads | 11 868 744 | 100 | 13 006 860 | 100 | 13 624 495 | 100 | 13 000 906 | 100 | 10 788 895 | 100 | 14 283 442 | 100 |
| no_adapter_reads | 657 906 | 6 | 851 701 | 7 | 630 224 | 5 | 1 034 491 | 8 | 767 790 | 7 | 689 372 | 5 |
| too_short_reads | 1 920 935 | 16 | 2 027 500 | 16 | 2 344 861 | 17 | 2 085 611 | 16 | 1 999 125 | 19 | 1 909 175 | 13 |
| UMI_defective_reads | 1 096 089 | 9 | 1 518 644 | 12 | 1 222 593 | 9 | 1 501 764 | 12 | 1 009 511 | 9 | 1 390 879 | 10 |
| Total reads - defaults | 8 193 814 | 69 | 8 609 015 | 66 | 9 426 817 | 69 | 8 379 040 | 64 | 7 012 469 | 65 | 10 294 016 | 72 |
| miRNA_Reads | 517 787 | 4 | 587 482 | 5 | 696 007 | 5 | 949 021 | 7 | 596 865 | 6 | 695 392 | 5 |
| hairpin_Reads | 13 336 | 0 | 15 300 | 0 | 12 592 | 0 | 16 669 | 0 | 10 947 | 0 | 16 699 | 0 |
| piRNA_Reads | 148 693 | 1 | 151 845 | 1 | 108 017 | 1 | 123 306 | 1 | 106 369 | 1 | 152 232 | 1 |
| rRNA_Reads | 113 033 | 1 | 161 104 | 1 | 138 852 | 1 | 129 807 | 1 | 92 205 | 1 | 121 951 | 1 |
| tRNA_Reads | 3 874 784 | 33 | 4 151 724 | 32 | 4 069 290 | 30 | 3 607 229 | 28 | 3 148 299 | 29 | 5 489 858 | 38 |
| mRNA_Reads | 192 852 | 2 | 188 884 | 1 | 266 619 | 2 | 187 639 | 1 | 133 068 | 1 | 176 971 | 1 |
| otherRNA_Reads | 811 183 | 7 | 829 882 | 6 | 1 167 023 | 9 | 1 294 313 | 10 | 846 707 | 8 | 1 143 937 | 8 |
| notCharacterized_Mappable | 706 579 | 6 | 743 437 | 6 | 839 704 | 6 | 654 517 | 5 | 620 655 | 6 | 753 382 | 5 |
| notCharacterized_notMappable | 1 815 567 | 15 | 1 779 357 | 14 | 2 128 713 | 16 | 1 416 539 | 11 | 1 457 354 | 14 | 1 743 594 | 12 |

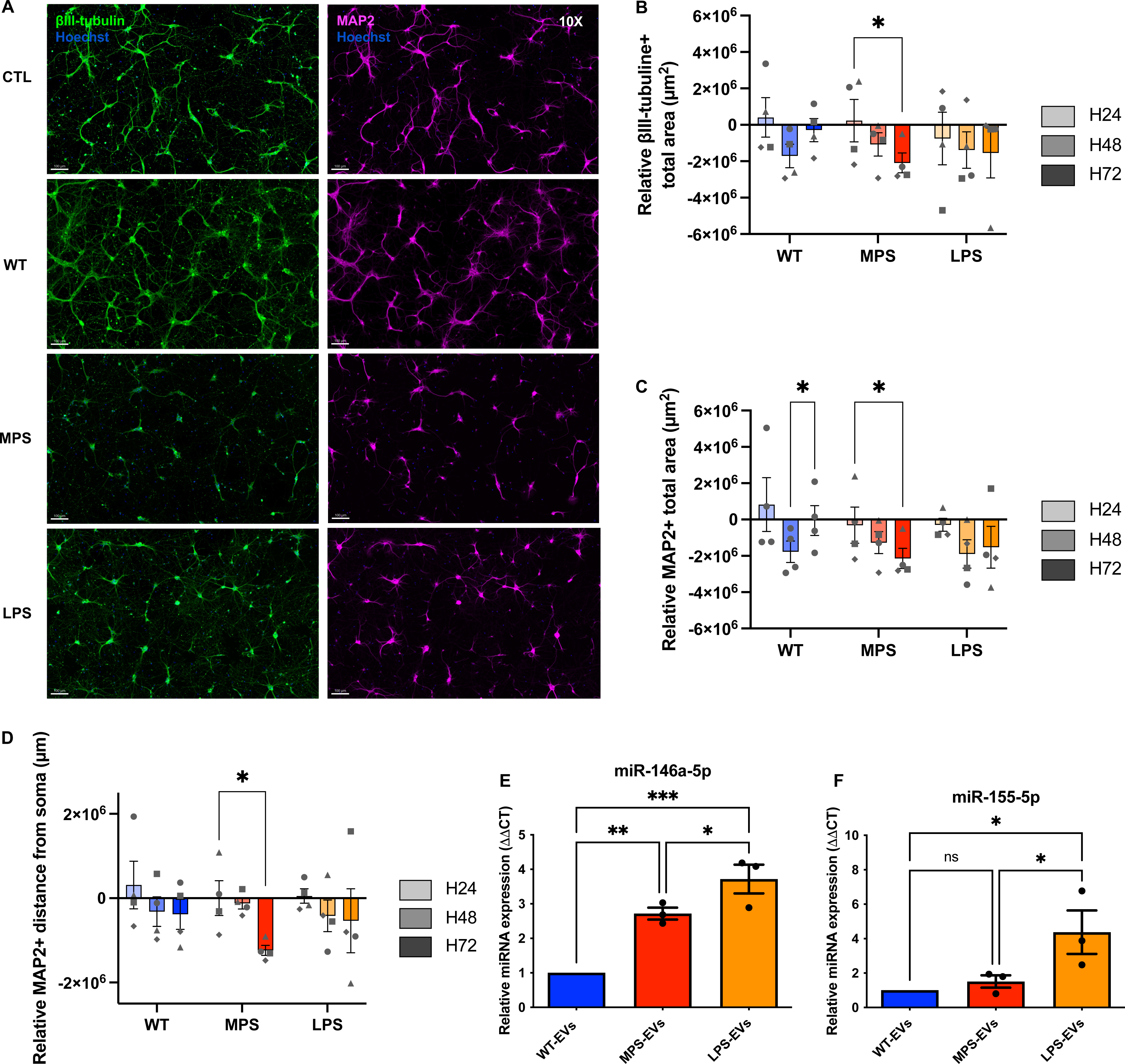
Data not shown: βIII-tubulin and MAP2 immunostaining of primary cortical neurons treated with WT-, MPS-EVs or LPS-EVs. **(A)** Representative insets of wide-field zoomed images of neurons on 25 squares mosaics (10X objective) showed a relative decrease in the intensity of MAP2 immunostaining in neurons after 72h of treatment with MPS-EVs or LPS-EVs. **(B-C)** Quantification of the total βIII-tubulin+ (B) and MAP2+ (C) surface area revealed a decrease in neurons treated with MPS-EVs but not with LPS-EVS or WT-EVs at 72h. **(D)** Quantification of the MAP2+ distance from soma revealed a decrease in neurons treated with MPS-EVs but not with LPS-EVS or WT-EVs at 72h. **(A-D)** Neurons were treated with WT-EVs (WT, blue), MPS-EVs (MPS, red), LPS-EVs (LPS, orange) or trehalose (CTL) for 24h (H24, in light grey), 48h (H48, in medium grey) or 72h (H72, in dark grey). Neurons were stained with an anti-βIII-tubulin antibody coupled to Alexa FluorTM 647 (green) or with an anti-MAP2 antibody coupled to Alexa FluorTM 568 (in pink). Nuclei were stained with Hoechst 33342 (in blue). Scale bar = 100 µm **(B, C and D)** Quantifications of wide-field neurons variables, normalized against non-treated control neurons (CTL). Two-way ANOVA, n=4 biological replicates (each point represents 1 n), * = p ≤ 0.05. **(E and F)** RT-qPCRs of miR-146a-5p (E) and miR-155-5p (F) represented by bar plots. RNA was extracted from EVs collected from conditioned media of BV-2 cells treated with WT-GAGs (WT-, blue), MPS-GAGs (MPS-, red) or LPS (LPS-, orange). One-way ANOVA, n=3 biological replicates (each point represents 1 n), ns = p > 0.05; * = p ≤ 0.05; ** = p ≤ 0.01; *** = p ≤ 0.001.

